# Integration of 168,000 samples reveals global patterns of the human gut microbiome

**DOI:** 10.1101/2023.10.11.560955

**Authors:** Richard J. Abdill, Samantha P. Graham, Vincent Rubinetti, Frank W. Albert, Casey S. Greene, Sean Davis, Ran Blekhman

## Abstract

Understanding the factors that shape variation in the human microbiome is a major goal of research in biology. While other genomics fields have used large, pre-compiled compendia to extract systematic insights requiring otherwise impractical sample sizes, there has been no comparable resource for the 16S rRNA sequencing data commonly used to quantify microbiome composition. To help close this gap, we have assembled a set of 168,484 publicly available human gut microbiome samples, processed with a single pipeline and combined into the largest unified microbiome dataset to date. We use this resource, which is freely available at microbiomap.org, to shed light on global variation in the human gut microbiome. We find that Firmicutes, particularly Bacilli and Clostridia, are almost universally present in the human gut. At the same time, the relative abundance of the 65 most common microbial genera differ between at least two world regions. We also show that gut microbiomes in undersampled world regions, such as Central and Southern Asia, differ significantly from the more thoroughly characterized microbiomes of Europe and Northern America. Moreover, humans in these overlooked regions likely harbor hundreds of taxa that have not yet been discovered due to this undersampling, highlighting the need for diversity in microbiome studies. We anticipate that this new compendium can serve the community and enable advanced applied and methodological research.

## Introduction

The human microbiome is an important factor in understanding health and disease. Systematic differences are observed between the composition of the microbiome in healthy individuals and those with microbiota-linked conditions such as colorectal cancer^1–3^ and inflammatory bowel disease.^4^ Thus, understanding and quantifying the determinants of variation in the microbiome has been a major goal of microbiome research. Studies have shown that this variation is driven by a variety of factors, including host genetics^5^ and ethnicity.^6–8^ While it is difficult to account for each of these factors individually, many are tied to geographic region. For example, dietary fiber and the consumption of processed foods varies between countries,^9^ as does the use of antibiotics,^10^ both of which are known to impact gut microbiota. Microbiome composition links location, culture and human health, a dynamic that can be observed in the compositional shifts experienced by individuals immigrating to the United States from Thailand,^11,12^ Latin America and Korea.^13^

Despite the importance of understanding microbiome variation between world regions, cultures, and social groups,^14–17^ many populations are practically excluded from the microbiome literature: In our previous work, we demonstrated that high-income countries, such as the United States, are dramatically overrepresented in public databases, while others, such as countries in eastern Asia, are under-sampled compared to their population.^18^ As with genome-wide association studies,^19^ a limited range of subjects raises the question of how broadly we can apply the known links between the microbiome and human health.^20,21^ The large number of publicly available microbiome datasets could be useful in quantifying differences between the most thoroughly studied world regions and those that are still comparatively uncharacterized.

In an environment as noisy and complex as the human gut, gaps in knowledge may be difficult to detect, and important patterns may only become apparent after collecting thousands or tens of thousands of samples. Large compendia such as ReCount^22,23^ that have been developed for transcriptomic analysis have revealed strain-level differences in complex microbial gene expression patterns^24^ and human gene expression modules that can be used to enhance transcriptome-wide association studies.^25^ The human microbiome field does not have a comparable resource.

To mitigate this gap in the bioinformatic capabilities of the field, we present here the Human Microbiome Compendium, a novel collection of more than 168,000 publicly available human gut microbiome samples from 68 countries. All samples were reprocessed using state-of-the-art tools and combined into a single dataset that we have made available in multiple formats, including the MicroBioMap R Bioconductor package and a website at microbiomap.org. We use this data to evaluate patterns in microbiome composition around the world and estimate the implications of gaps in our current knowledge of the human gut.

## Results

### Bacilli, Clostridia are universal constituents of the global human gut microbiome

To generate the Human Microbiome Compendium, we started with metadata for 245,627 samples of 16S amplicon sequencing available in the BioSample database maintained by the U.S. National Center for Biotechnology Information (NCBI) as of October 2021, limited to those submitted in the “human gut metagenome” category and flagged with the “amplicon” library strategy. The samples are organized into studies, as defined in the BioProject database; we processed each study separately using a pipeline centered around the DADA2 software tool,^26^ which generates a “taxonomic table” for each BioProject in which each row is a sample and each column is a single taxon. We elected conventional quality-control settings that should apply to the broadest number of studies: removing reads shorter than 20 nucleotides, for example, and reads with any ambiguous (“N”) base calls (see **Methods** for a comprehensive description of the pipeline and quality control). Briefly, studies for which paired-end reads could not be reliably merged were reprocessed as single-end after discarding reverse reads. We removed BioProjects with an elevated proportion of suspected chimeric reads, those for which taxonomic classification failed (indicating reads were not generated by conventional amplicon sequencing), and BioProjects for which the most abundant taxa were not bacterial, generally in studies focused on archaea or fungi. We focused on Illumina-based assays and discarded BioProjects reporting instruments that perform pyrosequencing or long-read sequencing.

To integrate the data across BioProjects, we processed and quantified each BioProject’s amplicon sequence variants (ASVs), each representing a single unique sequence observed in the samples in the BioProject. Then, each ASV was classified as specifically as possible down to the genus level. The final results quantified the number of reads in each sample that were assigned to each taxon. We repeated this for the 482 BioProjects in the complete dataset, which resulted in a full compendium of 168,484 samples from 68 nations, encompassing 5.57 terabases of sequencing data processed using a uniform pipeline (**Figure 1A**). Finally, for further analyses, we created a filtered compendium of 150,721 samples containing at least 10,000 reads each after excluding rare taxa (see **Methods**), to filter out low-quality samples with insufficient data on composition and microbes that are too rare to compare between BioProjects or world regions. The processed data, the Human Microbiome Compendium, is freely available in multiple formats at microbiomap.org (see **Data Availability**), where users can browse, visualize, filter, and download the data and metadata. We also created an R package, microbiomap, to facilitate further analysis of the Human Microbiome Compendium data (see **Data Availability**). Full documentation for the dataset and R package, as well as tutorials and examples of how to integrate the data into one’s own work, is available at microbiomap.org.

**Figure 1.**
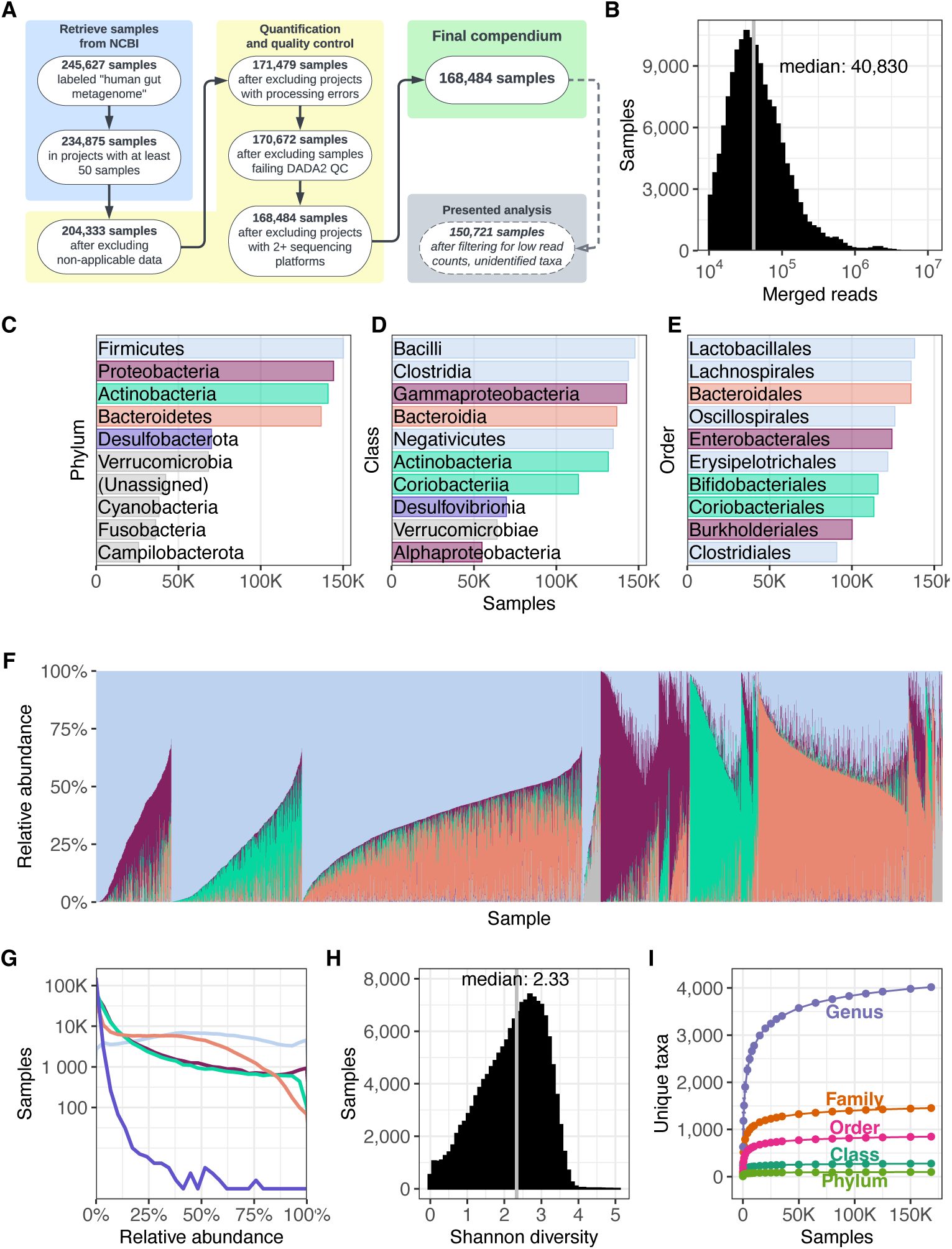
Overview of the Human Microbiome Compendium. **(A)** A list of the general steps in the data pipeline and how many samples completed each step. See **Methods** for more details about each process. **(B)** A histogram illustrating the distribution of reads that were classified in each sample. The x-axis indicates the number of reads in a given sample, and the y-axis indicates the number of samples with that number of reads. **(C–E)** The most prevalent taxa observed in the compendium. The reads in each sample are assigned the most specific taxonomic name possible, down to the genus level. Each panel illustrates results when these assignments are consolidated at the three highest taxonomic levels; in each, the y-axis lists the 10 most prevalent taxa at that level, and the x-axis indicates the number of samples in which that taxon was observed at any level. Panel **C** indicates the most prevalent phyla, and the top five are each assigned a color. These colors are used in the remaining two panels to indicate the phylum of each taxon. Panel **D** indicates the most prevalent classes of bacteria observed in the dataset, and Panel **E** indicates the most prevalent orders. Lower taxonomic orders are illustrated in Supplementary Figure 1. **(F)** A stacked bar plot illustrating the relative abundance of 5000 randomly selected samples from the compendium. Each vertical bar represents a single sample, and the colored sections each represent the relative abundance of a single phylum in that sample. These bars use the same colors as **panel C**. The samples are sorted first by the most abundant phylum’s identity, followed by the second-most abundant phylum’s identity, followed by the combined relative abundance of these two taxa. For example, the first group on the left is made up of samples in which Firmicutes was the most abundant phylum and Proteobacteria was the second-most abundant. Next is samples in which Firmicutes was most abundant and Actinobacteria was second-most prevalent, and so on. Another version of this figure, sorted by Firmicutes relative abundance, is available as Supplementary Figure 2. **(G)** A density plot illustrating the relative abundance of phyla across the compendium. Each line represents one of the five most prevalent phyla in the dataset, using the same colors as **panel B**. The gray line indicates all other phyla. The x-axis indicates the relative abundance of a given phylum in a single sample, and the y-axis indicates how many samples were observed to have that abundance of the given taxon. A version of this figure using a linear y-axis is available as Supplementary Figure 3. **(H)** A histogram illustrating the distribution of Shannon diversity observed in the compendium. The x-axis indicates a given sample’s alpha diversity, as measured by Shannon Diversity Index. The y-axis indicates the number of samples that were observed to have that score. **(I)** The results of a rarefaction analysis in which a simulated compendium of various sizes was generated repeatedly and evaluated for taxonomic richness. The x-axis indicates the number of microbiome samples in the simulated compendium, and the y-axis indicates the number of unique taxa were observed in that simulation. Each line indicates the number of observed taxa at successively specific taxonomic levels.

The median sample contained 40,830 reads, after trimming, quality filtering, and merging of paired reads, and 90.4 percent of samples had fewer than 150,000 reads (**Figure 1B**). As observed even in early sequencing assays of the human gut microbiome,^27^ we find the Firmicutes phylum is by far the most prevalent (Supplementary Figure 1), found in 150,540 of 150,721 samples (99.9 percent), followed by Proteobacteria (144,489 samples; 95.9%), Actinobacteria (141,191 samples; 93.7%) and Bacteroidetes (136,984 samples; 90.9%), before a sharp drop-off to phyla such as Desulfobacterota and Verrucomicrobia (**Figure 1C**). Firmicutes also contains three of the five most prevalent classes (**Figure 1D**) and five of the 10 most prevalent orders (Supplementary Table 1). The prevalence of the Bacteroidetes phylum, long a focus of analysis (e.g. ref^28^) is due almost entirely to the Bacteroidales order in our data (136,085 samples; 99.3 percent of Bacteroidetes-positive samples; **Figure 1E**), particularly the Bacteroidaceae family.

Visual inspection of the phylum-level relative abundances found in the compendium shows a surprisingly uniform distribution for the abundance of Firmicutes (**Figure 1G**), which is in the top two phyla of 137,091 samples (91.0 percent) and combines with Bacteroidetes, the fourth-most prevalent phylum, to make up the majority of the reads classified in 51.0 percent of samples (Supplementary Table 2). In samples with lower abundances of Firmicutes and Bacteroidetes, a limited number of phyla take their place (**Figure 1F**; Supplementary Figure 2): Of the 73,859 samples in which at least one other phylum appears in the top two, Actinobacteria is a top-two phylum in 52.1 percent of them, followed by Proteobacteria (46.2 percent). Of the 4,750 samples for which neither Firmicutes nor Bacteroidetes is in the top two, Proteobacteria is in the top two of 97.5 percent. We find the relative abundance distribution of Proteobacteria closely resembles that of Actinobacteria, and that Desulfobacterota, despite being the fifth-most prevalent phylum by appearing in 70,302 samples, is found at relative abundances lower than 1 percent in 88.8 percent of those samples (**Figure 1G**; Supplementary Figure 3).

We observed a wide range of alpha diversity (a measure of taxonomic richness within samples), with a median Shannon diversity of 2.33 and values as high as 5.07 (**Figure 1H**), consistent with ranges identified in previous meta-analyses of alpha diversity across multiple microbiome studies.^29,30^ To estimate the completeness of this census, we performed a sample-based rarefaction analysis,^31^ in which we selected random subsamples of different sizes without replacement from the full compendium and evaluated the number of unique taxa observed in each subsample (see **Methods** for details). The discovery rate for new taxa approached zero after 25,000 samples for all levels except genus, the most specific. Between subsamples of 150,000 samples and the full dataset of 168,484, we observed one new genus every 4831 samples (**Figure 1I**). This demonstrates that the compendium currently captures all but the rarest taxa in the populations covered by the dataset, given the current distribution of reads per sample. Overall, we find that there is broad variation in the composition of human gut samples (**Figure 1F**), but within a limited selection of microbial taxa drawn mostly from the Firmicutes phylum (**Figure 1E**).

### World regions harbor unique microbiome signatures

Though much of the public metadata available for BioSamples is inconsistently reported ^32^, the “geo_loc_name” identifier was available for 92.4 percent of samples in the filtered compendium. This field provides general information about where a sample was collected, and in most cases directly specifies the country of origin. We manually reviewed all 455 unique “geo_loc_name” values (Supplementary Table 3) and associated each with a standardized list of countries (see **Methods**); we then consolidated all of these countries into eight world regions defined by the United Nations Sustainable Development Goals (SDG) program (**Figure 2A**). As in previous work,^18^ we found that the majority of samples were from Europe and Northern America (91,144 samples; 60.5 percent), with the Eastern and South-Eastern Asia region a distant second at 17,086 samples (11.3 percent; **Figure 2B**). Sub-Saharan Africa was the third-most represented (5538 samples; 3.7 percent), followed closely by Central and Southern Asia (5046 samples; 3.4 percent).

**Figure 2.**
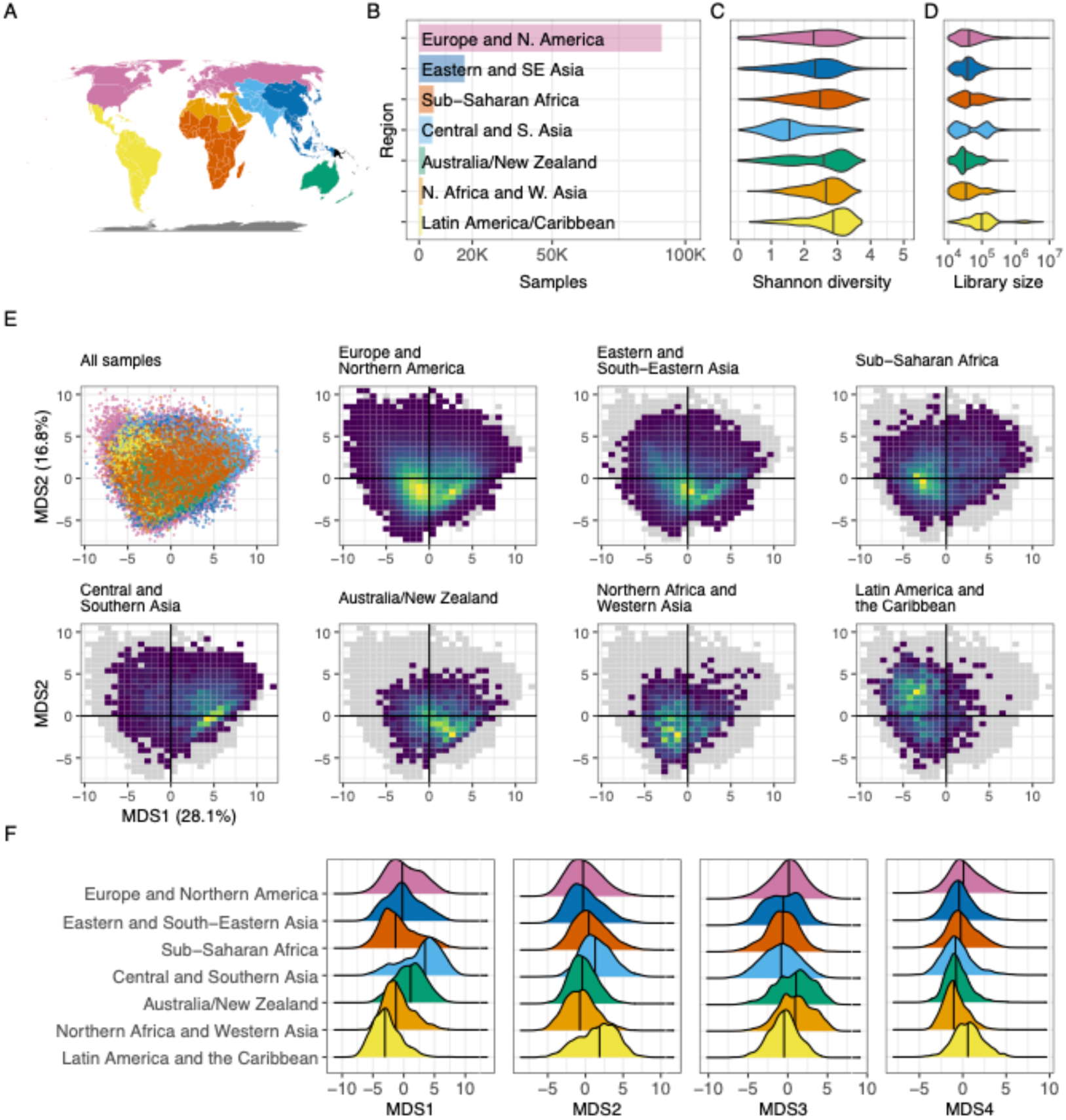
Regional structure. **(A)** A map illustrating which areas were categorized into world regions. The colors here match those labeled in panel **B**. Oceania is represented here in orange, though this region was excluded from these analyses because only four Oceanis samples remained in the filtered dataset used here. **(B)** A bar plot illustrating the number of samples from each world region analyzed here. The x-axis illustrates total samples, and the y-axis lists all regions evaluated. The colors used here are the same as those used in panel **A**. **(C)** A violin plot illustrating the distribution of observed Shannon index values assigned to samples from each world region. The x-axis indicates the Shannon index value, as calculated using all unique taxonomic identifications in samples from each world region. Colors indicate the region (same as in **A**), and the y-axis for each violin indicates the relative frequency with which diversity of a given magnitude was observed. The vertical lines in each violin indicate the median value. The black points within each violin indicate the mean Shannon diversity as determined by rarefaction analysis (see **Methods**). **(D)** A violin plot organized in the same manner as panel **C**, but the x-axis indicates reads per sample. “Reads” in this case refers to merged reads that were included in the filtered taxonomic table. **(E)** A series of plots illustrating the results of a principal coordinates analysis of samples from all world regions. The top-left plot is a scatter plot in which each point is a single sample; the color indicates the sample’s region, using the scheme described in panel **A**. The x-axis is the first PCoA axis, which explains the most variation across the dataset; the y-axis is the PCoA axis explaining the second-most variation. The seven other plots use the same axes, but each includes only samples from a single world region. These plots use a heatmap design rather than a scatter plot, to help evaluate areas with many overlapping points—yellow areas indicate portions of the space with a higher concentration of samples, and dark blue areas indicate portions in which few (but not zero) samples are found. The gray shadow indicates the area occupied by all points from all world regions. **(F)** A series of density plots illustrating the distributions of the first four axes of variation determined by the ordination analysis displayed in panel **E**. Each panel illustrates a single factor; the x-axis indicates the value of that factor, and the y-axis indicates the relative frequency of the value in the given world region.

We observed the highest alpha diversity among the 1195 samples from Latin America and the Caribbean, with a median Shannon index of 2.90 (**Figure 2C**). Samples from Central and Southern Asia exhibited the lowest average diversity (median Shannon index=1.55), though it’s possible we are underestimating the diversity of under-studied world regions because of gaps in reference databases used for taxonomic assignment (see **Discussion**). Pairwise comparisons show significant differences in alpha diversity between all world regions (Wilcoxon rank-sum test, q<10^-4^; Supplementary Table 4) except for Australia/New Zealand, which was not significantly different from Sub-Saharan Africa (q=0.61). We also reconsidered this analysis because of differences in read depth between regions: The Australia/New Zealand region had the lowest median reads per sample (30,415 reads; **Figure 2D**), compared to the average of 98,641 reads from the most deeply sequenced samples of Latin America and the Caribbean. To account for this, we performed a rarefaction analysis (see **Methods**) that allowed us to determine an average Shannon diversity while controlling for both reads per sample and samples per region (Supplementary Figure 5); the mean Shannon diversity was reduced slightly in all regions, as expected because of lower read counts, but none of the means differed by more than 0.2 percent (Supplementary Table 5).

To explore differences in microbiome composition across world regions, we used principal coordinates analysis (PCoA) for visualization of the dataset (**Figure 2E**; see **Methods**). We plotted the samples from each region on the same axes calculated for the dataset as a whole, allowing for direct comparison between groups along two dimensions that together account for 44.9 percent of variation (Supplementary Figure 6). Though this approach meant the analysis was heavily weighted in favor of variance observed in Europe and Northern America, using these axes helps to illustrate whether samples from the rest of the world differ from those of the most thoroughly characterized region. Even in these two dimensions, we observe systematic differences between world regions—some, such as Australia/New Zealand, appear to cluster in subsets of the main areas occupied by Europe and Northern America, but others, such as Latin America and the Caribbean, occupy areas of the projected space that are far less commonly observed elsewhere. This pattern continues when we evaluate more than the first two axes of variation: We can observe large-scale regional differences in the distributions of the first four axes (**Figure 2F**), and, using all eight axes extracted from the dataset (Supplementary Figures 7–13), clusters defined by world region are much more compact and distinct than would be expected by random (Davies–Bouldin index=6.93; p<4×10^-6^; Supplementary Figure 22). Though there is substantial overlap between the world regions, unintuitive differences are apparent in the first two axes. For example, the “hot spots” of Central and Southern Asia occupy a different space in the ordination plots than the samples of Latin America and the Caribbean (**Figure 2E**), while the samples of Europe and Northern America occupy a very similar area to the samples from Eastern and South-Eastern Asia. Together, these results demonstrate a link between microbiome composition and geography, even at this high level in which some regions encompass many countries home to billions of people.

### Uneven microbiome sampling leaves taxa undiscovered

To investigate microbiome diversity in world regions, we repeatedly subsampled each region and identified the number of unique microbial taxa present in the selected microbiome samples (**Figure 3A**; Supplementary Figures 14–17). For this analysis, we used all 4,018 taxa quantified by the initial DADA2 assignment and all samples with a world region assignment. Notably, more taxa are discovered in samples from Eastern and South-Eastern Asia than any other region, despite having around 70,000 samples fewer than the largest region. Approximately 2.5 percent of samples from Eastern and South-Eastern Asia contain more than 300 distinct taxa, but this many taxa are found in only 0.05 percent of samples from Europe and Northern America. 3,662 of the 4,018 taxa are present in samples from Eastern and South-Eastern Asia; only 3,299 and 874 of these taxa are present in Europe and Northern America and Latin America and the Caribbean, respectively.

**Figure 3.**
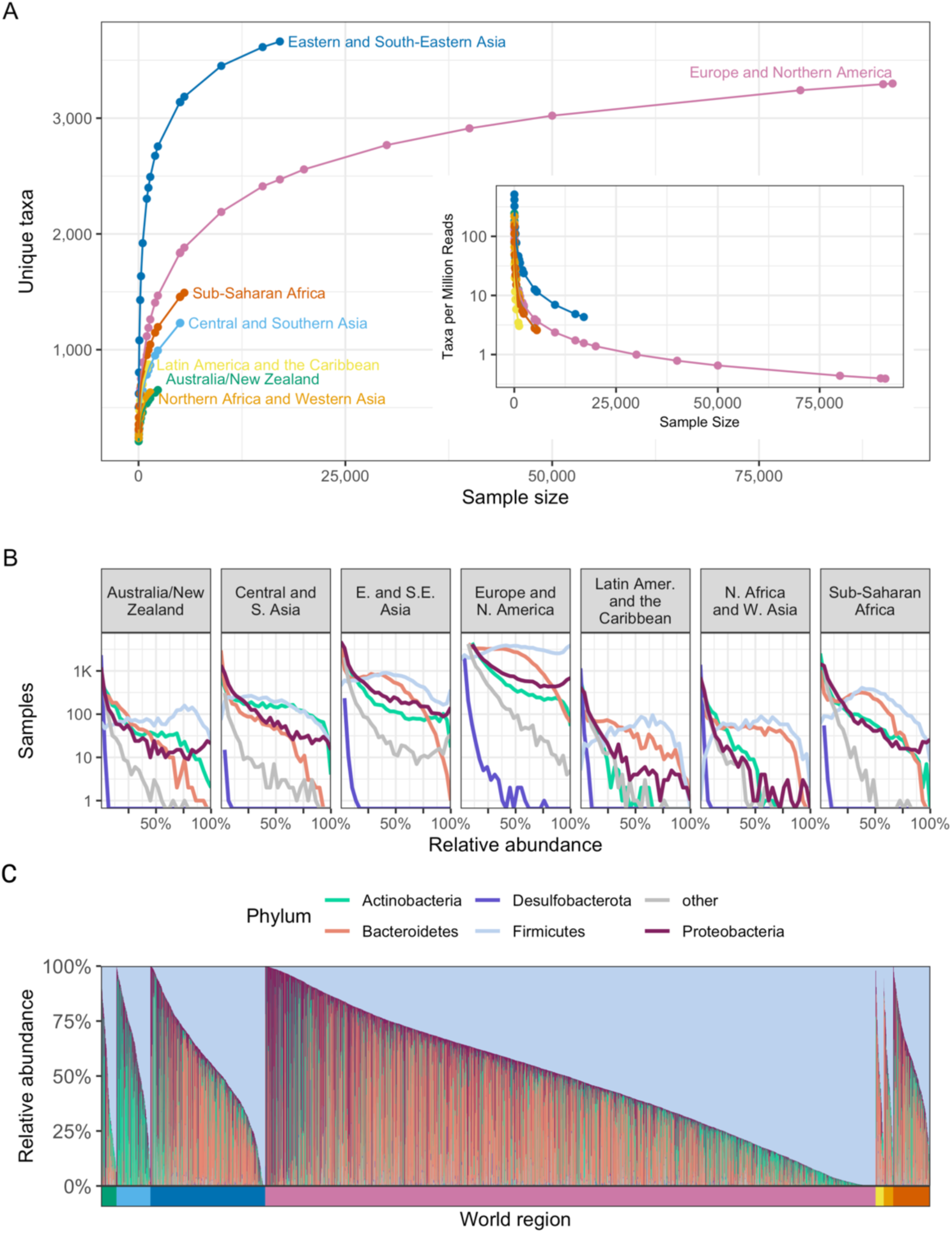
Geographic regions vary in microbiome composition. **(A)** The number of unique taxa discovered in subsamples of varying size from each world region. Each point represents the average number of unique taxa identified in a subsample from a given region over 1,000 repetitions. The x-axis indicates the number of microbiome samples selected, the y-axis the number of unique taxa identified in those samples, and the color indicates the world region being sampled. The inset uses the same x-axis and color scheme but displays the average number of taxa discovered per million reads on the y-axis. **(B)** Histograms illustrating the distribution of the relative abundance of the most prevalent phyla in the compendium. Each panel visualizes all samples from a single world region. The x-axis indicates the relative abundance of the taxon, and the y-axis indicates the number of samples (on a log scale) with the indicated relative abundance. Each line illustrates the results for a single phylum, indicated by line color. **(C)** As in Figure 1F, this stacked bar chart shows the relative abundance of the five most prevalent phyla in the compendium. Each column is a sample, and the colored segments indicate the relative abundance of a given phylum in that sample. Phylum color follows the same color scheme as Figure 3B. Samples are ordered first by world region (indicated by the colored bar below the x-axis), and then by relative abundance of the 5 most prevalent phyla, as in Figure 1F. World region color follows the same color scheme as Figure 3A.

We note that the rate of discovery drops for each world region after the first few thousand samples. On average, 371 more taxa were discovered in 2,000 samples from Eastern and South-Eastern Asia than in 1,000 samples, yet only 210 more taxa were found in 17,086 samples (all samples from the region) than in 10,000 samples in the same region, demonstrating the decline in discovery rate. This decline in discovery rate holds true for the most sampled region as well: 2,190 unique taxa were identified in the first 10,000 samples from Europe and Northern America, but the subsequent 81,144 samples uncovered only 1,109 new taxa.

The rate of discovery for Europe and Northern America falls below one new taxon per million reads by the time 30,000 samples are assayed (**Figure 3A inset**), indicating the sequencing effort required to identify new taxa in this region: when all samples from this region are included, the discovery rate is a mere 0.39 taxa per million reads. By comparison, when including all samples from Northern African and Western Asia, we continue to discover an average of 8.19 taxa per million reads (Supplementary Figure 16). In Latin America and the Caribbean, the discovery rate is 3.03 new taxa per million reads when all samples from the region are assayed. In fact, other than Europe and Northern America, the lowest discovery rate observed is 2.58 taxa per million reads, when all samples are assayed in Sub-Saharan Africa. The stark difference in discovery rate between Europe and Northern America and that of the next-lowest region emphasizes the unequal sampling between world regions, and the relatively high discovery rate in world regions other than Europe and Northern America indicates that many taxa may be uncovered with further sampling in underrepresented regions.

While the top phyla remain consistent across world regions, their abundance and prevalence differ (**Figure 3B**; Supplementary Figures 18–19): In Europe and Northern America, the relative abundance of Firmicutes is approximately uniformly distributed. In Sub-Saharan Africa, however, the distribution of Firmicutes peaks at approximately 40 percent, with higher relative abundances becoming less and less common. Samples in Northern Africa and Western Asia have nearly identical distributions of Firmicutes and Bacteroidetes, though we do observe broad patterns of diversity within world regions as well (**Figure 3C**). While Firmicutes is dominant in each region, samples from Central and Southern Asia have higher relative abundances of Actinobacteria (Wilcoxon test, p<2.2×10^-16^ for Central and Southern Asia vs other) than the other regions, plus correspondingly lower abundances of Bacteroides (Wilcoxon test, p<2.2×10^-16^ for Central and Southern Asia vs other; **Figure 3C**).

### Systematic differences in microbiome composition between world regions

To quantify how specific microbial taxa vary between world regions, we performed differential abundance analysis using a linear mixed model that accounts for BioProject, mean ASV length, and amplicon used for sequencing (Supplementary Table 6), as these experimental artifacts may bias results (see **Methods**). We focused our analysis on the 65 genera that had a minimum of 1 percent prevalence and 0.5 percent relative abundance in at least one world region (see Methods). Pairwise comparison of each region revealed distinct differences among these genera, and all 65 taxa tested were found to be significantly differentially abundant between at least one pair of regions (Supplementary Table 7). The highest number of differentially abundant taxa (56) were found when comparing samples from Sub-Saharan Africa to samples from Australia/New Zealand, while the fewest differentially abundant taxa (8) were found between Europe and Northern America and Northern Africa and Western Asia (**Figure 4A**).

**Figure 4.**
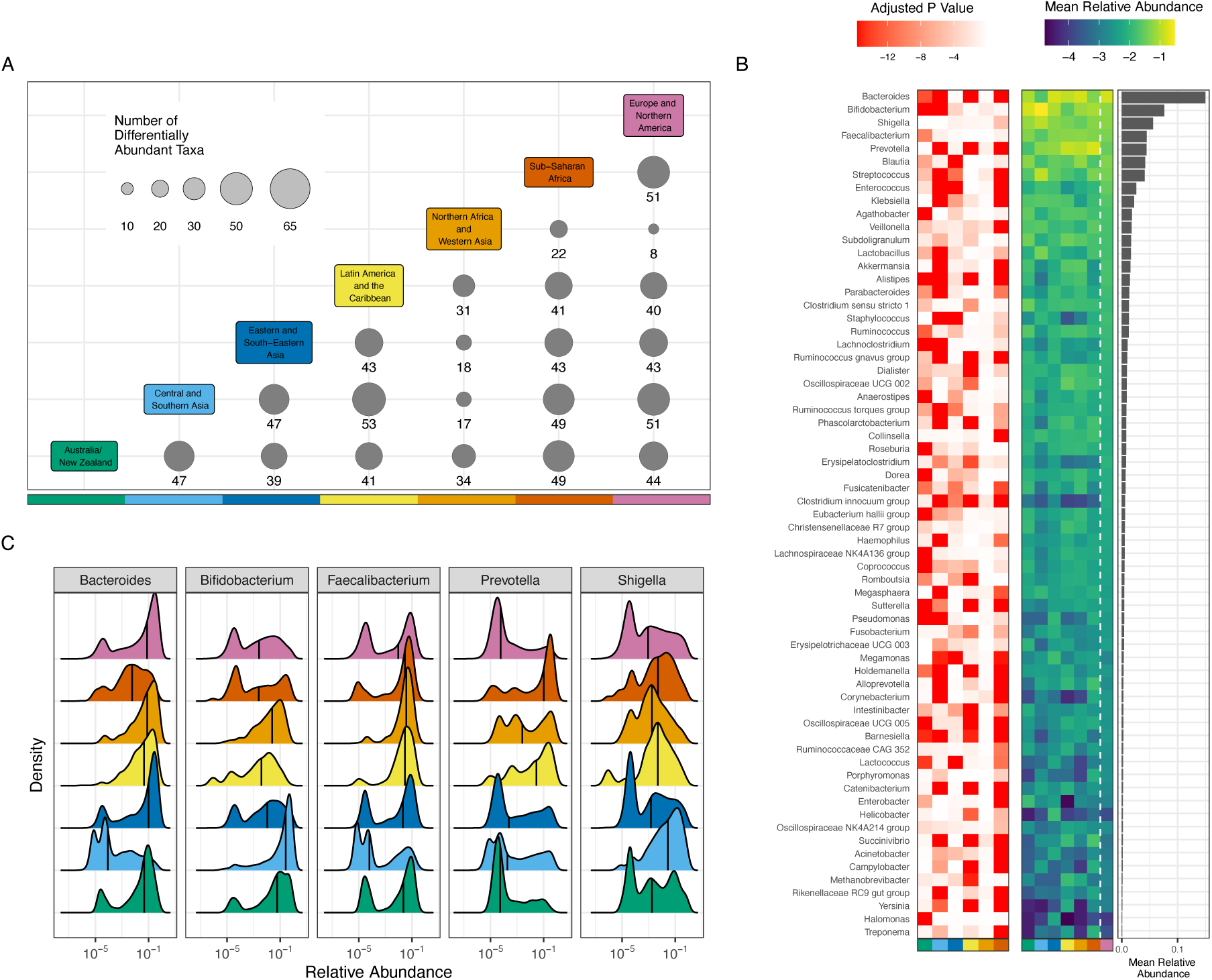
Taxa are differentially abundant between world regions. **(A)** 65 taxa were selected to be tested for differential abundance between regions. The x and y axes are each colored by world region; at each intersection, the size of the circle and the number underneath it indicate the number of taxa that were significantly different between the two regions listed. **(B)** The red-white heat map illustrates adjusted p-values for regional differences when each world region is compared to Europe and Northern America. The y-axis lists all evaluated genera, the x-axis lists each region (using the same color scale as panel **A**), and each cell represents the strength of the differential abundance result for that taxon. The blue-green heat map illustrates mean relative abundance (log 10) of each taxon in each world region, as indicated by the x-axis. The bar chart illustrates the mean relative abundance of each taxon across all regions. **(C)** Each panel illustrates the relative abundance (log 10) of one of the 5 most abundant taxa. Each colored area indicates the distribution from a single world region, using the same colors as panel **A**. The x-axis indicates (log 10) relative abundance of the specified genus, and the y-axis indicates the relative frequency with which that abundance is observed in the specified region. Black vertical lines indicate the median.

As samples from Europe and Northern America make up over half of the compendium, subsequent analyses focused on the differences found between Europe and Northern America and each of the other world regions, to evaluate the most broadly observed differences relative to the most sampled region (Supplementary Figures 20–21). Prevotella and Bacteroides were two genera with the lowest adjusted p-value between Europe and Northern America and any other region, indicating strong differences between world regions (**Figure 4B).** Bacteroides abundance is higher in Europe and Northern America than Sub-Saharan Africa (q<2.2×10_-16_), Latin America and the Caribbean (q<2.2×10^-16^), Central and Southern Asia (q<2.2×10^-16^), and Australia/New Zealand (q=7.87×10^-14^). Conversely, Prevotella abundance is lower in Europe and Northern America than Sub-Saharan Africa (q<2.2×10^-16^), Latin America and the Caribbean (q<2.2×10^-16^), and Central and Southern Asia (q<2.2×10^-16^). Concordant with prior literature,^33^ we observe a higher relative abundance of Bacteroides in Europe and Northern America than in non-westernized regions such as Sub-Saharan Africa and Central and Southern Asia, and we observe an increase in the abundance of Prevotella in Sub-Saharan Africa as compared to Europe and Northern America (**Figure 4B).**

Closer examination of the distribution of relative abundances of different genera across world regions reveals more specific patterns. Nearly every region has a high number of samples with high abundance of Bacteroides, as evidenced by the strong peak close to 1 in **Figure 4C**. Notably, Central and Southern Asia and Sub-Saharan Africa appear to have fewer samples with such high relative abundance of Bacteroides, as these regions lack a strong peak at 1 in **Figure 4C**. Over 26 percent of samples from Europe and Northern America contain more than 30 percent Bacteroides; only 4.3 percent and 2.2 percent of samples from Sub-Saharan Africa and Central and Southern Asia, respectively, contain as much Bacteroides. Prevotella, a taxon commonly associated with positive health outcomes and non-western microbiomes, appears to be higher in abundance in Sub-Saharan Africa q<2.2×10^-16^) and Latin America and the Caribbean (q<2.2×10^-16^) when compared to Europe and Northern America. Only 19 percent of samples from Europe and Northern America have more than 1 percent Prevotella–visible in the strong peak near 0 in **Figure 4C**–compared to 43 percent of samples from Northern Africa and Western Asia, 61 percent of Latin America and the Caribbean, and 65 percent of Sub-Saharan African samples.

After evaluating compositional differences between regions, we then sought to define region-specific signatures of gut microbiomes by identifying the taxa most closely linked to the overall variance observed in principal components analysis, performed separately for each region (termed here the ‘variance score’; see **Methods**). We find variability in Europe and Northern America is best represented by the relative abundances of Escherichia/Shigella (variance score=0.98; Supplementary Table 8), Enterococcus (0.97), Lactobacillus (0.95), Akkermansia (0.94) and Bifidobacterium (0.93), while the microbiomes of Northern Africa and Western Asia are defined by the genera Prevotella (0.85), Shigella (0.81), Akkermansia (0.77), Dialister (0.57) and Bacteroides (0.52). Of the top five taxa in each regional signature, all are members of the six most prevalent phyla in the compendium (**Figure 1C**). Escherichia/Shigella was the only genus to appear in the top 5 taxa for all evaluated regions. Using the top 10 taxa in each regional signature, we found that six genera appear in the signatures of all world regions: Bifidobacterium, Bacteroides, Prevotella, Streptococcus, Veillonella, and Shigella, which together form what we could consider the core taxa most useful for explaining global variation in the human gut microbiome—not necessarily the most prevalent, but the taxa that vary most widely in all regions. Oppositely, there are five genera that appear in the top 10 taxa for only a single region: Staphylococcus (Central and Southern Asia), Megamonas (Eastern and South-Eastern Asia), Dialister (Northern Africa and Western Asia), Collinsella (Sub-Saharan Africa), and Alistipes (Latin America and the Caribbean).

## Discussion

Here, we integrated data from 168,484 publicly available 16S rRNA amplicon sequencing samples from 482 BioProjects to evaluate global variation in the human gut microbiome. We found the majority of available samples were from Europe and Northern America, which has been so extensively sampled that most microbiota present in the region’s gut microbiomes have likely already been observed, while further samples from other regions may uncover up to 20 times as many new taxa per million reads. Thousands of unique taxa have also been observed in Eastern and South-Eastern Asia, but samples show such remarkable diversity that there are likely many more yet to be uncovered. Though practically all taxa are shared to some degree between world regions, we found each region occupies a unique niche within the ordination space defined via multidimensional scaling, identified dozens of taxa that are differentially abundant between each region, and determined strong regional signatures indicating the primary gradients that define the composition of microbiomes around the world.

Others have articulated the vital importance of studying microbiomes from diverse populations^20,21^ and evaluating the potential consequences of inaction.^15,34^ Though our analysis uses only samples from previous works, compiling hundreds of disparate studies enabled the evaluation of differences we can observe even given the comparatively limited sample sizes. We find that variance in Europe and Northern America, by far the most thoroughly sampled region, is closely tied to the relative abundance of Lactobacillus (variance score=0.95), which has been linked to obesity in the United States^35^ and bipolar disorder in Austria.^36^ It remains to be answered how these results should be interpreted in Latin America and the Caribbean, where Lactobacillus is practically absent from the regional signature (variance score=0.03) though not necessarily absent from the microbiomes there. We also found many taxa with consistent differences in world regions when compared to Europe and Northern America (**Figure 4B**), including highly abundant genera such as Bacteroides, Bifidobacterium and Prevotella, the abundance and proportions of which may play a role in inflammation,^37^ obesity,^38^ inflammatory bowel disease,^39^ and the early development of the pediatric gut microbiome,^40^ among many other conditions.

Still, there is reason for caution in drawing strong conclusions from such a broad range of samples collected for very different reasons in hundreds of projects. First, world region may be confounded with why the samples were collected, and the data currently does not have consistent metadata related to host health. Relatedly, reference databases may have less coverage of taxa that appear more commonly outside of Europe and North America,^41,42^ which would result in more unidentified taxa and deflated diversity estimates in samples from other regions of the world. Regarding our analysis, any combination of studies raises concerns about batch effects, or artifactual findings that are caused by technical details but appear to be of biological origin.^43,44^ We are optimistic that these effects are minimized in large-scale analyses—in our previous work, we found batch correction of large compendia is ineffective when there are many “batches” (in this case, projects)—as the number of batches grows, the disparate project-level effects are overshadowed by the legitimate biological signal, which is more consistent across studies.^45^ Lastly, the dataset compiled for this project does not resolve the broad issue of representational imbalances in global human microbiome research,^18,46,47^ though there are many ongoing projects that seek to increase the diversity of microbiome research—projects such as the African Microbiome Program are expanding work not only in humans, but agricultural microbiomes as well,^48^ and initiatives like H3ABioNet aim to address some of the structural challenges to expanding the populations under study.^49,50^

In addition to our findings on global variation, we are also optimistic about the Human Microbiome Compendium’s utility as a way to better utilize existing resources: The National Institutes of Health have directly invested more than $1 billion in human microbiome research,^51^ and raw data for tens of thousands of microbiome samples are uploaded to SRA every year, plus many more from collaborators in the International Nucleotide Sequence Database Collaboration (INSDC), which includes organizations in Japan and the European Union. Although these are world-class repositories for a huge variety of genomic data, this raw data is difficult to manage at scale: The primary utility of compendia such as recount3, a database of uniformly processed RNA-seq data, is that these sequencing reads have already been processed, curated and combined together into a unified dataset. There are several microbiome resources like this: the MicrobiomeHD project integrated data from 28 case–control studies ^52^; the most recent version of GMrepo includes 45,111 amplicon sequencing samples ^53^; and the curatedMetagenomicData project^54^ now includes data from 22,588 whole-metagenome shotgun samples.^55^ These projects focus on human-curated data with uniform metadata, a valuable asset to the field. However, these projects still represent a small fraction of the available samples; we hope our compendium, several times larger than those currently available, will be useful in situations where sample size is a more important factor than thorough annotation, such as performing further meta-analysis or providing context for other datasets (e.g. ref^56^).

In summary, we present here the Human Microbiome Compendium, a new, large-scale collection of human gut microbiome data. We use this compendium to study microbiome variation at a global scale, comparing world regions and showing that some regions likely have many taxa that remain undiscovered due to undersampling. We expect this compendium will be a valuable resource for the community and enable novel insights into the microbial ecology of the human gut.

## Methods

### Sample selection

We retrieved metadata for all BioSamples categorized in the NCBI Taxonomy^57^ under “human gut metagenome” on 9 October 2021. After removing samples that could not be associated with a BioProject or sequencing run, we selected only those for which the library source was “genomic” or “metagenomic,” excluding the values “metatranscriptomic,” “transcriptomic,” “viral RNA,” “synthetic” and “other.” Of these, we then limited the dataset to samples with a “library strategy” value of “amplicon” (and not values such as “WGS” and “RNA-Seq”). This left 245,627 samples across 1,437 BioProjects. We then excluded BioProjects that contained less than 50 samples meeting our criteria, leaving us with a list of 234,875 samples in 811 BioProjects.

Because pyrosequencing technologies developed by companies such as 454 Life Sciences and Ion Torrent require different processing steps, we then sought to remove BioProjects containing pyrosequencing data and other sequencing instruments that use processes dissimilar from Illumina sequencing, such as MinION. We used the SRA Toolkit APIs to retrieve sequencing instrument information for each sample and evaluated BioProjects that reported using 454 or Ion Torrent instruments. We found 19 such instruments (“454 GS FLX Titanium,” “Ion Torrent PGM,” “454 GS FLX,” etc.), but manual review revealed one instrument was consistently mislabeled: In many BioProjects (PRJNA685914 and PRJNA605031, for example) the sequencing instrument was reported as “454 GS” even though the authors report elsewhere in the BioProject that they used sequencers such as Illumina’s MiSeq platform, which we were not attempting to remove. More careful examination revealed these BioProjects (and many others) performed their analysis using Mothur,^58^ a popular microbiome analysis tool, and “454 GS” is Mothur’s default entry for the “instrument” field when uploading to SRA.^59^ To avoid removing applicable samples, we removed BioProjects reporting using any of the pyrosequencing instruments except for “454 GS.”

### Sequencing data retrieval

Samples were exported from the database one BioProject at a time; each BioProject had a file listing all accessions of runs associated with samples meeting the criteria described above. This file was used as the input for the “fasterq-dump” tool^60^ from the SRA Toolkit maintained by NCBI. This tool downloads the data from the Sequence Read Archive and splits the information into FASTQ files for downstream processing. We used samples from all available INSDC members—of the completed samples, 126,452 were from the Sequence Read Archive (74 percent), 38,971 were from European Nucleotide Archive, and 5249 were from the DNA Data Bank of Japan. We did not include projects in which more than 10 samples failed to download, which was generally caused by files missing from SRA.

### Amplicon processing

If the number of files for forward reads matched the number of files for reverse reads, we processed the BioProject as paired-end sequencing. If there was a mismatch, or there were no reverse reads, we processed the BioProject as single-end data. In both cases, we used DADA2 v1.14.0 to process the data.^26^ We used general settings that we believed would be effective across many BioProjects,^61^ aiming to maintain as many samples as possible while excluding low-quality data: We did not trim a set number of bases from either end, nor did we limit the maximum length of a read. We removed reads shorter than 20 nucleotides, reads with any ambiguous (“N”) base calls, and any reads that aligned to the phiX genome (if present, almost certainly as a control in Illumina sequencing runs). We also disabled quality-based truncation of reads. Paired-end reads were merged with a minimum overlap of 20 bases. In some cases, the process of merging reads failed, and close to zero forward reads were merged with their paired reverse read, likely due to sequencing strategies that involve non-overlapping reads or reads with very minimal overlap. For BioProjects in which fewer than 50 percent of forward reads were merged successfully, we discarded the reverse reads rather than concatenating them, to avoid situations in which merging failed because of low-quality calls or mismatched forward and reverse read files. In those cases, the reverse reads were removed, and these BioProjects were re-processed as single-end data. If the number of forward reads did not match the number of reverse reads in a sample, we attempted to use DADA2 to detect the sequence identifier field in the FASTQ file to match the samples that could be salvaged. If this was unsuccessful, we removed the reverse reads and reprocessed these as single-end data as well. Taxonomic assignment was performed by DADA2 using the SILVA database release 138.1.^62,63^ We believe this was the most reliable way to process this data, given the lack of information about project-level sequencing strategies. The merging process in paired-end datasets would be much more effective with more knowledge of study design and primer choices, for example, particularly in cases where the amplicon length was greater than the read length and the paired-end reads did not overlap. In addition, DADA2 recommends building separate error models for each sequencing run,^64^ but only BioProject could be reliably inferred, which means ASV inference at the study level may not capture run-level patterns.

Though we removed obviously non-applicable samples (mycobiome assays using the 18S rRNA gene rather than 16S, for example), we did not pursue more stringent filtering. To minimize the number of legitimate samples removed from the compendium, we avoided making discretionary decisions about removing samples or BioProjects from the compendium that we encountered during analysis—for example, 199 samples from BioProject PRJEB25853 are included because they were deposited in the “human gut metagenome” category, though manual review of the metadata shows the samples are actually from a novel assay of the vaginal microbiome.^65^ We plan to expand the automated curation processes used to review results for future iterations of the compendium but will continue to favor approaches that are too permissive rather than risk filtering out legitimate samples with compositions that don’t “look right.” We also plan to integrate more metadata from both samples and projects to enable more precise filtering.

### Pipeline success

Due to resource constraints, we did not attempt to process BioProjects with fewer than 50 samples, which accounted for 10,752 out of 245,627 samples. Of the 234,875 samples we processed, we found 31,887 samples (13.6 percent) contained non-applicable data— BioProjects that targeted fungi or archaea, for example, and mislabeled BioProjects that used shotgun or nanopore sequencing instead of 16S amplicon sequencing. Another 31,509 samples (13.4 percent) were excluded because extracting acceptable results, if possible at all, would have required manual intervention and more knowledge of the sequencing strategy: DADA2 identified excessive chimeric reads in some BioProjects, for example, and we excluded any BioProject in which at least five of the first 10 samples processed contained more than 25 percent chimeric reads. Several BioProjects were also excluded because they contained samples associated with multiple sequencing runs, or were associated with samples that could not be downloaded. 807 samples were removed from BioProjects because all of their reads were filtered out.

### Dataset for analysis

We combined BioProject-level taxonomy tables into one large matrix containing 168,464 samples (from 482 BioProjects) for rows and 4018 unique taxonomic identifiers for columns, making up the Human Microbiome Compendium. For most of the analysis reported in this paper, we applied additional quality control steps: First, we removed 16,781 samples (10 percent) with fewer than 10,000 reads. We then removed 2018 taxonomic entries (50 percent) with fewer than 1000 total reads across all remaining samples, and a further 578 taxa (14 percent of the original) that were detected in fewer than 100 samples. After these taxa were removed, we again evaluated the sample read counts and removed another 19 samples that now had less than 10,000 reads. (The removal of these 19 samples did not push any more taxa below the above thresholds.) We then evaluated the proportion of reads in each sample for which a taxonomic assignment could not be assigned at at least the phylum level. We removed 943 samples for which more than 10 percent of the sample’s reads had an unassigned phylum. This left us with 150,721 samples and 1422 taxa. This filtered compendium was used in almost all presented analysis, except for Figures 1I and 3A where we used the unfiltered compendium with 168,464 samples.

### Calculation of Shannon diversity

To calculate the Shannon index for individual samples, we used the “diversity” function in the vegan R package v2.6.4,^66,67^ using natural logarithms and all columns in the filtered dataset—that is, ASVs were consolidated if their taxonomic assignments matched exactly, but counts were not consolidated at any single taxonomic level. This data was also used to calculate each sample’s Simpson’s Index and species count via vegan’s “diversity” and “specnumber” functions respectively.

### Rarefaction analysis, taxon discovery rate

We estimated the relationship between compendium size (in number of samples) and total taxa observed by performing a sample-based assessment.^31^ Specifically, we built simulated compendia of various sizes between 1 and 150,000 by subsampling the filtered analysis dataset and counting the number of unique taxa observed in each subsample at each taxonomic level. We repeated each compendium size simulation 150 times and plotted the mean observed taxa at each taxonomic level; this allowed us to build curves plotting observed taxa against compendium size.^68,69^ Generally, the x-axis of this curve would plot *total reads*, rather than *total samples*, to account for variation in the number of “observations” (i.e. a single read from a single microbe) in different size compendia. Here, we visualized total samples instead (**Figure 1H**) to incorporate the differences in read depth actually observed in the data. The trade-off is that these metrics will likely underestimate *future* taxa observed if used to extrapolate forward into larger compendium sizes, as the distribution of reads per sample (**Figure 1B**) will likely shift as time passes and sequencing costs continue to drop.

### World region alpha diversity comparison

We compared alpha diversity measurements between all regions using the Wilcoxon rank-sum test,^70^ with multiple test correction done using Hochberg’s method^71^ as implemented in the R “stats” package’s “wilcox.text” and “p.adjust” functions respectively. The calculations were performed using the filtered compendium dataset used in other analyses, but was limited to samples with a world region annotation other than “unknown.”

### Rarefaction analysis, regional alpha diversity

We estimated regional alpha diversity (in the filtered analysis dataset) by selecting 1000 random samples from each world region (enough that each region could provide all samples without replacement), then rarefied each of these samples down to 10,000 randomly selected reads each. From these rarefied samples, we determined the mean Shannon diversity for each region, then repeated the entire process 1000 times.^72^

### Country and region inference

We were able to obtain a “geo_loc_name” tag from the NCBI BioSample database (https://www.ncbi.nlm.nih.gov/biosample/) for 155,584 of 168,484 samples (92.4 percent). These tags held 455 unique values, which were associated with countries by manually reviewing the values and associating them with a country (Supplementary Table 3). Most tags contained enough context to confidently assign a country name: The most common tag value was “usa:new york” found in 14,142 samples from six BioProjects, followed by “usa” (12,510 samples in 46 BioProjects). However, the third-most common tag was “missing” (5532 samples; 27 BioProjects), and other tags such as “not applicable” (4317 samples) and “not available” (3481 samples) appeared many times. Overall, we were able to assign a country to 447 of the 455 unique values (98.2 percent) representing 153,152 samples (90.9 percent). In total, we found 68 countries represented in the geo_loc_name values—to simplify comparisons, we consolidated these assignments into eight world regions defined by the United Nations Sustainable Development Goals (SDG) program (**Figure 2A**).

### World region inference accuracy estimate

To assess the accuracy of our process for associating samples with their country (and therefore region) of origin, we selected a random sample of microbiomes and manually determined the country of origin (Supplementary Table 9), primarily by finding publications referencing the data but also using other metadata associated with the samples and parent BioProjects. We found the “geo_loc_name” tags to reliably include country names, which gave us confidence in our ability to infer country from tag, but the additional factor of interest is whether samples were mislabeled by the original authors. Between these two factors, we assumed 95 percent accuracy for our sample-size calculation, which was designed to detect this level of accuracy at a precision of ±5 percent at a 95 percent confidence interval.^73^ This results in a sample size of 73.0; after accounting for a 25 percent “dropout” rate (samples with a “geo_loc_name” tag but no other means with which to verify it), our new sample size was 97.3. Because some BioProjects are 60 times larger than others, we wanted to mitigate the effect of selecting many samples from one large, single-country BioProject by first selecting 100 random BioProjects, then selecting one sample from each project to evaluate. To verify the accuracy of our inferences, we first looked for an explicit statement of the study’s country of origin in the project description in the BioProject database. If this did not yield an answer, we looked to any publications linked to the BioProject and searching Google Scholar for several factors indicating a link to the BioProject: First, we searched the BioProject accession, then its corresponding SRA accession (such as “ERP006059” for BioProject PRJEB6518), then its ID number (such as “bioproject 589558” for BioProject PRJDB6499), then any unique phrases from the BioProject title or description. If the paper did not explicitly state a country of origin for the subjects, we considered the classification confirmed if the paper included ethical approval from an institutional review board in that country. If none of these steps could confirm a country, we classified it as a dropout. Of 100 samples evaluated, eight were dropouts. Two were deemed not applicable because the papers described samples that were incubated prior to sequencing. Of the remaining 90, we were able to validate that all 90 had world region assignments that were confirmed by either the BioProject description or a publication associated with the data. With a 100 percent success rate, we can then use the rule of three^74^ to estimate that the lower bound of the confidence interval (at our original 95 percent confidence level) is 96.67 percent accuracy.

### Principal coordinates analysis

We began by using the matrix of read counts to build a distance matrix between all samples using the robust Aitchison distance.^75^ We then performed multidimensional scaling (**Figure 4**) using the “divide and conquer” approach described by Delicado and Pachon-Garcia^76^ and implemented in the “bigmds” R package. We extracted 8 principal coordinates and used 16 points as the overlap between partitions.^77^

### World region cluster evaluation

We measured the effectiveness of regional clustering by calculating the Davies–Bouldin Index^78,79^ of the regional clusters formed in the 8-dimensional space generated by the ordination described above. This gave us a single score (6.93), but no context for comparison, so we estimated a p-value by performing a bootstrap analysis in which we generated 250,000 additional scores using the same data but with regional labels that were shuffled without replacement. This provided a distribution to which we could compare the real score, but the observed range of values in the shuffled data was 83.98–468.73, resulting in an estimated p-value of 0.

### Differential abundance analysis

We further filtered the dataset used for analysis for genera with a mean relative abundance of at least 0.5 percent and prevalence of at least 1 percent in any world region, resulting in 65 remaining genera. We then filtered the samples to include only those with at least 1000 reads in the 65 genera. This resulted in analysis of 123,346 samples and 65 genera. To test these taxa for differential abundance, we used a linear mixed model using the lme4 and lmerTest R packages.^80,81^ We modeled world region as a fixed effect, and to account for technical artifacts included BioProject, mean ASV length, and amplicon as random effects. We ran a single model for each taxon, running each model 7 times so that each world region could be the reference variable, to enable pairwise comparison between each region–region pair as follows:

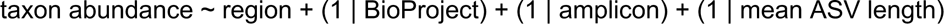

To account for the inherent compositionality of microbiome data, the taxon abundances were centered log-ratio transformed after adding a pseudocount of 1 to any 0 values, and subsequently scaled to a mean of 0 and standard deviation of 1. The p-values outputted from each model were multiple-test corrected using Benjamini-Hochberg. For figure presentation, p-values lower than 2.2×10^-16^ were adjusted to 2.2×10^-16^. The full results of this analysis are reported in Supplementary Table 7.

### World region taxon discovery rate

To generate the region-level taxon discovery rate curves shown in **Figure 3A**, we performed the following analysis: For each world region, we selected *n* random microbiome samples from the region and recorded the number of unique taxa present in these samples. We repeated this subsampling 1,000 times for each sample size, and reported the mean in Figure 3A. We repeated this strategy for all listed sample sizes in all world regions. This data is available in Supplementary Table 10. To verify that the curves for each world region in **Figure 3A** are distinct, we performed pairwise Wilcoxon rank-sum tests on each region-region pair for all possible sample sizes, comparing the number of unique taxa discovered over all 1000 repetitions for each region. This data is available in Supplementary Table 11. The inset panel in **Figure 3A** was generated using the same sampling strategy, but we also recorded the total number of reads in the selected samples. We then calculated the number of taxa discovered per one million reads for each repetition, and reported the mean in the inset in Figure 3A. We calculated 95 percent confidence intervals for this value by bootstrapping with 1000 repetitions and present this data in Supplementary Figure 16.

### World region signature determination

We used principal components analysis (PCA) to extract the taxa that are most closely linked to overall variance observed in the microbiomes of each region, which results in a heuristic we refer to here as the variance score. This score ranges from 0, indicating no relationship, to 1, indicating that the major axes of variation in the region also perfectly explain the variation observed in that taxon. We started with the taxonomic table of each region, applied the robust centered log ratio transformation to the read counts,^75^ then applied PCA to each region separately. We kept as many principal components (PCs) as was required to account for at least 50 percent of variance in each region’s data. We then used the resulting eigenvectors to calculate a score for each taxon observed in that region that indicates how much variance of that taxon was explained by the selected PCs. These proportions range between 0 and 1 and are used as the variance score for each taxon. The key assumption is that if a subset of principal components explains the majority of variance in the dataset, and those same PCs explain a high proportion of variance for a single taxon, then that taxon is more strongly linked to overall variability in the dataset than taxa that are poorly captured by the selected PCs.

### Amplicon inference

We used the amplicon sequence variants (ASVs) generated by DADA2 for each BioProject to infer the sequencing strategy used by each BioProject—primarily determining which of the nine hypervariable regions were targeted for amplification, but also the size of the amplicon targeted. To do this for each BioProject, we retrieved the sequence of all ASVs detected in that BioProject. We aligned each ASV to the sequence of the full *E. coli* 16S rRNA gene sequence, obtained from GenBank^82^ under accession J01859.1,^83^ using the striped Smith– Waterman library^84^ integrated into scikit-bio v0.5.8.^85^ If the optimal alignment covered 70 percent or less of the full ASV sequence, the alignment was discarded and the ASV was classified as unknown. For the remaining ASVs, the coordinates of the alignment were used to determine which of the nine hypervariable regions were covered by the ASV, as defined by Chakravorty et al.^86^ A region was considered to be covered if more than half of its length was covered by the ASV—for example, if an ASV’s alignment starts just before the beginning of V3 and ends 60 bases into the 107-base V4 region, that ASV would be classified as “V3–V4.” If the same alignment ended only 20 bases into the V4 region, that ASV would be classified only as “V3.” Beginning region and ending region (i.e. “V3” and “V4” from this example) were tallied separately. If more than half of all ASVs were categorized in a single starting region, that region was determined to be the starting region for the entire BioProject. If more than half of all ASVs were categorized in the same ending region, that region was determined to be the ending region for the entire BioProject. In situations where the threshold was met for only the starting or ending region (generally because of wide variation in ASV length), the opposite region was determined using the known region and the average ASV length. In situations where the ending region was determined to be before the starting region, the assignments were discarded under the assumption that this indicated multiple sets of primers were used.

### Website for high-level exploration

We designed a website to serve as an entry point to the Human Microbiome Compendium, hosted at https://microbiomap.org. The website displays important links to downloads and materials, provides a brief overview of the project and data, and features controls and visualizations for answering basic questions about the data. A search box allows users to check if a sample, BioProject, or taxon of interest is present in the data, and interactive maps and charts show sample counts at different geographic and taxonomic levels. Users can select a country or world region on the map, and the taxonomic level chart will adjust to show sample counts associated with the selected feature.

The website was implemented as a React single-page application, scaffolded and bundled with Vite, to allow for cleaner implementation of interactive features. The D3 library was used for more complex visualizations and interactions. The entire project utilizes TypeScript to ensure full static type safety.

The website’s data is generated by a set of pre-processing “compile” scripts, which automatically download the latest Human Microbiome Compendium data files from the place of record, then restructure and pare-down the data into a more practical, static subset for efficient display on the web. These scripts run before any build of the website, and can also be run on a schedule or on-demand.

### Bioconductor package implementation

We implemented a Bioconductor^87^ package, MicroBioMap (https://github.com/seandavi/MicroBioMap), that provides convenient access to compendium data. Data are loaded into a Bioconductor TreeSummarizedExperiment object,^88^ providing opportunities to use our compendium data with extensive Bioconductor microbiome analysis and visualization tools. The package includes documentation and example use cases.

### Software tools

The diagram in Figure 1A was created using Lucidchart (https://lucidchart.com). Most analyses were performed using R 4.2.2. Analyses requiring a high-performance computing environment, the rarefaction curves in Figure 1 and the multidimensional scaling analysis from figure 2, used R 4.3.1. Maps use the Equal Earth projection^89^ and the rnaturalearth R package.^90^

## Other information

### Supplementary information

**Supplementary Figure 1:** Prevalence at the family and genus levels.

**Supplementary Figure 2:** Relative abundance across samples.

**Supplementary Figure 3:** Relative abundance of top phyla.

**Supplementary Figure 4:** Diversity across world regions.

**Supplementary Figure 5:** Rarefaction diversity estimates.

**Supplementary Figure 6:** Scree plot for the compendium-wise ordination analysis.

**Supplementary Figures 7–12:** Ordination plots.

**Supplementary Figure 13:** Ordination results.

**Supplementary Figure 14:** Region-level discovery curve with 95% confidence intervals

**Supplementary Figure 15:** Region-level discovery curve (x-axis: sample size vs y-axis: taxa per read)

**Supplementary Figure 16:** Region-level discovery curve (x-axis: number of reads vs y-axis: unique taxa)

**Supplementary Figure 17:** Region-level discovery curve (x-axis: sample size vs y-axis: taxa per million reads) - same as 3A inset, but with error bars

**Supplementary Figure 18:** Histograms from fig 3B with individual y-axes

**Supplementary Figure 19:** Histograms from fig 3B with linear y-axes

**Supplementary Figure 20:** Ridgeline plots of differentially abundant taxa 6–35

**Supplementary Figure 21:** Ridgeline plots of differentially abundant taxa 36–65

**Supplementary Figure 22:** Cluster strength analysis.

**Supplementary Table 1:** Observed prevalence at each taxonomic level

**Supplementary Table 2**: Frequency of combinations of top-two phyla in each sample. Each row indicates the most abundant phylum in a sample, each column indicates the second-most abundant, and the cells indicate how many samples were observed with a given combination.

**Supplementary Table 3:** geo_loc_name values associated with countries

**Supplementary Table 4:** Pairwise world region alpha diversity comparison.

**Supplementary Table 5:** Regional rarefaction summary statistics

**Supplementary Table 6**: BioProject-level inferred metadata on amplicon choice and ASV length

**Supplementary Table 7:** Differential abundance analysis results

**Supplementary Table 8:** Regional signatures

**Supplementary Table 9**: World region inference analysis notes

**Supplementary Table 10:** Region-level discovery rate results

**Supplementary Table 11:** Results of Wilcoxon rank-sum test to compare region-level discovery rate curves

## Acknowledgements

We thank the members of the Blekhman lab for helpful discussion. We also thank Chad L. Myers, R. Stephanie Huang, Anna Selmecki, and William Harcombe for advice. This work was completed with resources provided by the University of Chicago’s Research Computing Center and the Minnesota Supercomputing Institute. This work was supported by NIH grant R01LM013863 (to R.B. and C.S.G.).

## Data availability

Data and code from this study is available on multiple platforms:

● The full microbiomap dataset is available for download from Zenodo at https://doi.org/10.5281/zenodo.8186993
● The open-source microbiomap R package is available on GitHub at https://github.com/seandavi/MicroBioMap
● The code used for data processing of the raw data, plus the code used to generate the figures in this manuscript, is available on GitHub at https://github.com/blekhmanlab/compendium_v1.
● An interactive website at **microbiomap.org** includes summary data for the compendium, links for downloads, and information about ongoing updates to the project.

## Author contributions

Conceptualization: RJA, SPG, RB, FWA

Data curation: RJA, SPG, VR

Formal analysis: RJA, SPG

Software: RJA, VR, SD

Supervision: FWA, CSG, SD, RB

Writing – original draft: RJA, SPG, VR

Writing – review & editing: FWA, CSG, SD, RB

