## Supplementary figures for "Integration of 168,000 samples reveals global patterns of the human gut microbiome"

Abdill, Graham et al.

**Supplementary Figures**

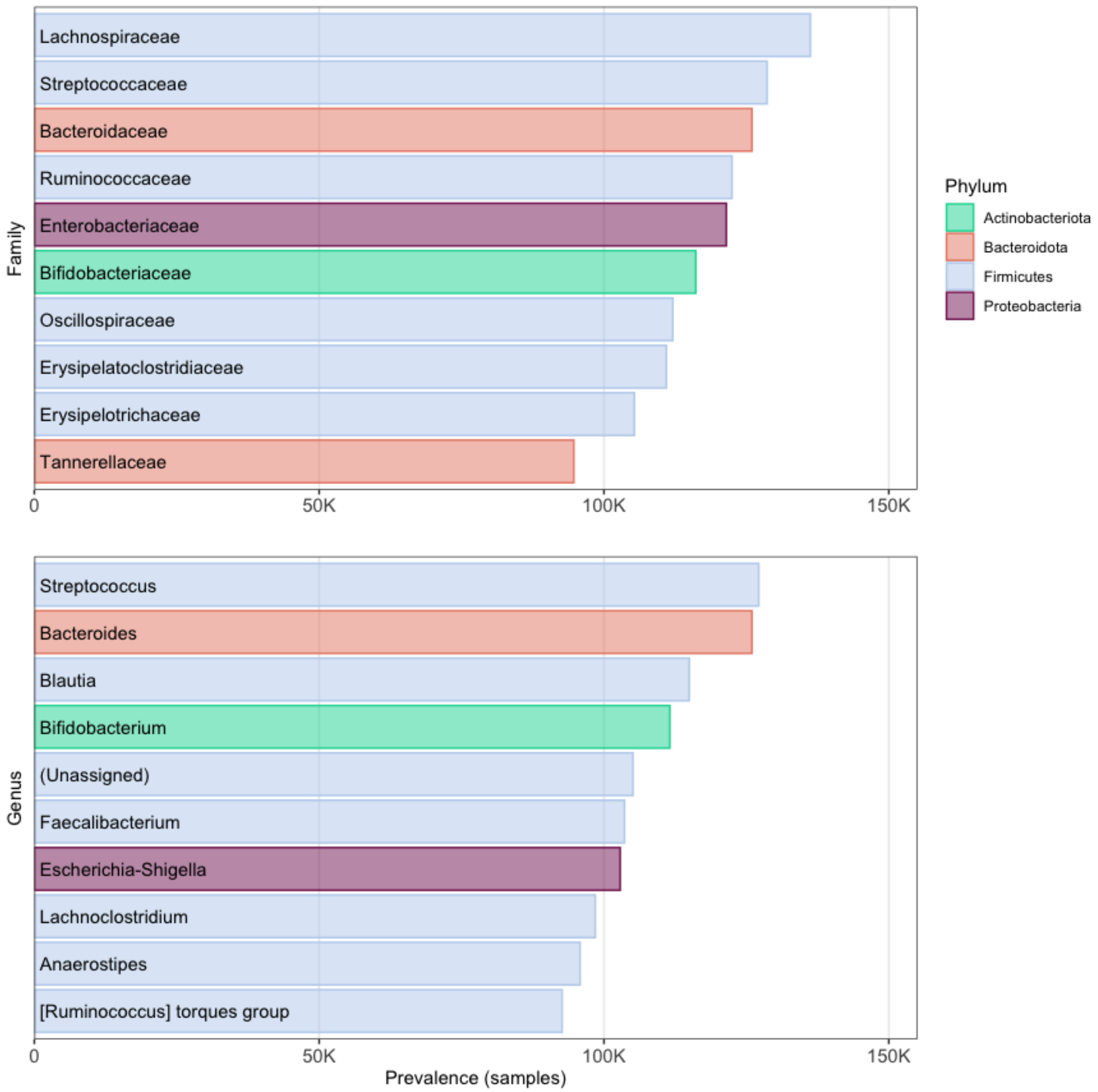


**Supplementary Figure 1. Prevalence at the family and genus levels.** The most prevalent taxa observed in the compendium. This figure extends the information in **Figure 1C–E**, which lists higher taxonomic orders. The reads in each sample are assigned the most specific taxonomic name possible, down to the genus level. Each panel illustrates results when these assignments are consolidated at the family level (top panel) and genus level (bottom); in each, the y-axis lists the 10 most prevalent taxa at that level, and the x-axis indicates the number of samples in which that taxon was observed. The five most prevalent taxa in the compendium are each assigned a color, which are used in the panels to indicate the phylum of each taxon.

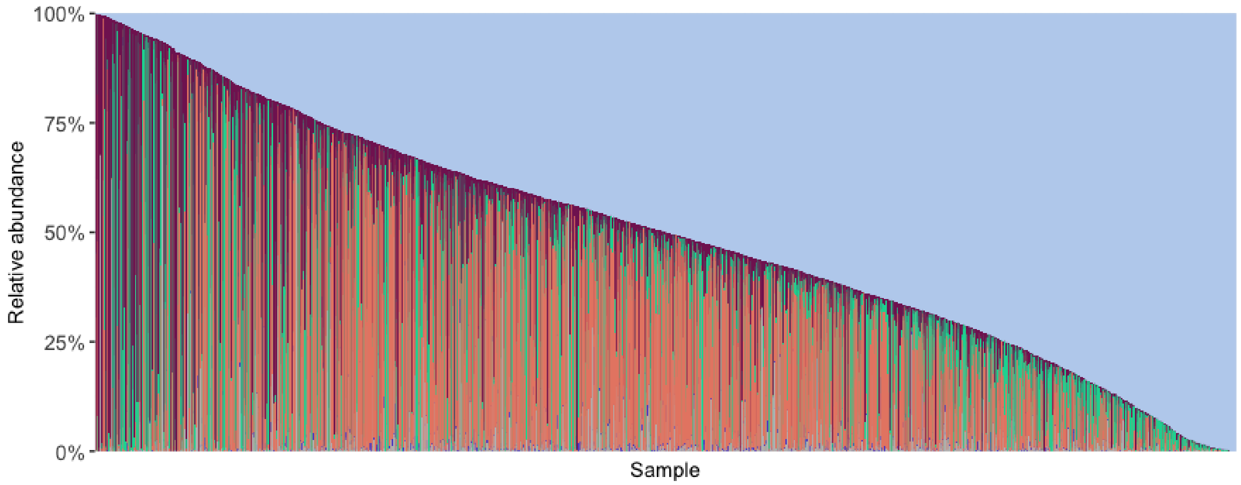


**Supplementary Figure 2. Relative abundance across samples.** Another version of the data illustrated in **Figure 1F**. A stacked bar plot illustrating the relative abundance of 5000 randomly selected samples from the compendium. Each vertical bar represents a single sample, and the colored sections each represent the relative abundance of a single phylum in that sample. Samples are ordered by the relative abundance of Firmicutes, the most prevalent phylum.

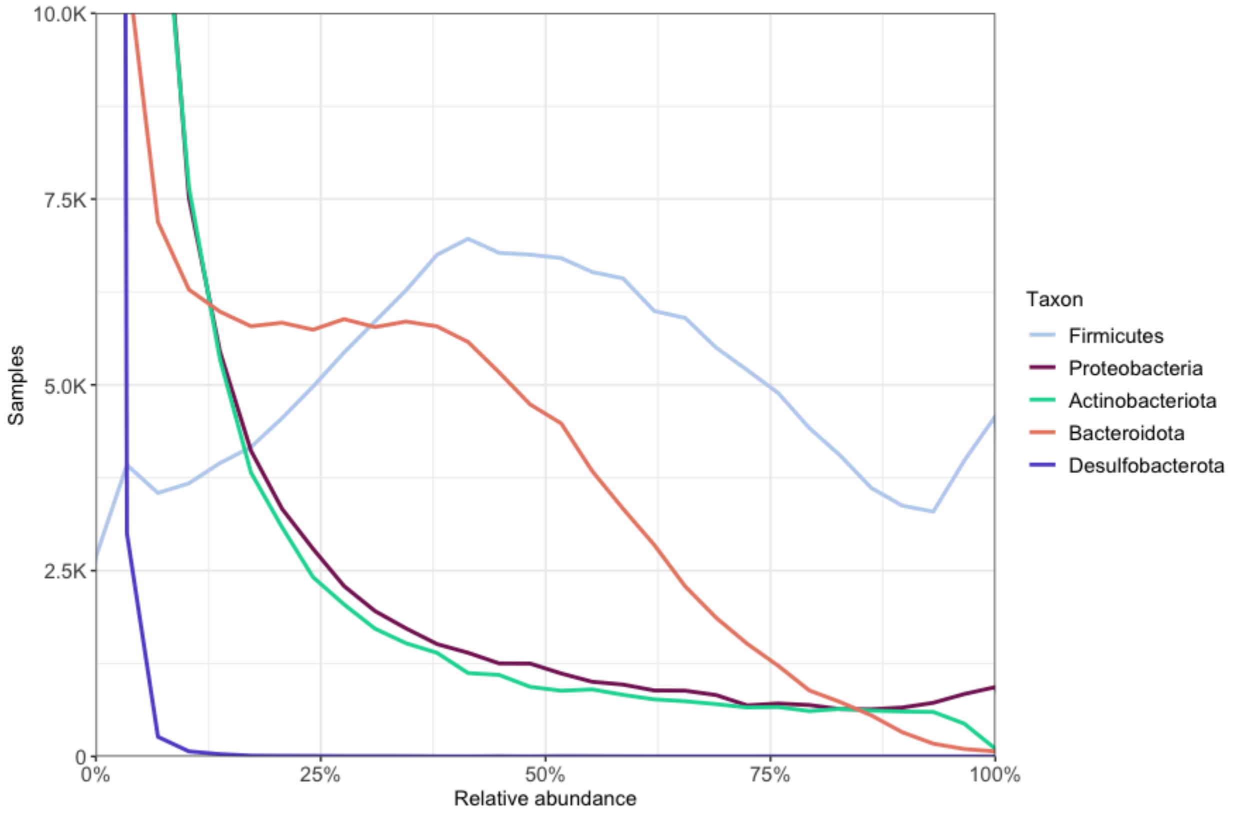


**Supplementary Figure 3. Relative abundance of top phyla.** This is another version of the data illustrated in **Figure 1G**, but using a linear y-axis that is truncated at 10,000 samples. A density plot illustrating the relative abundance of phyla across the compendium. Each line represents one of the five most prevalent phyla in the dataset. The x-axis indicates the relative abundance of a given phylum in a single sample, and the y-axis indicates how many samples were observed to have that abundance of the given taxon.

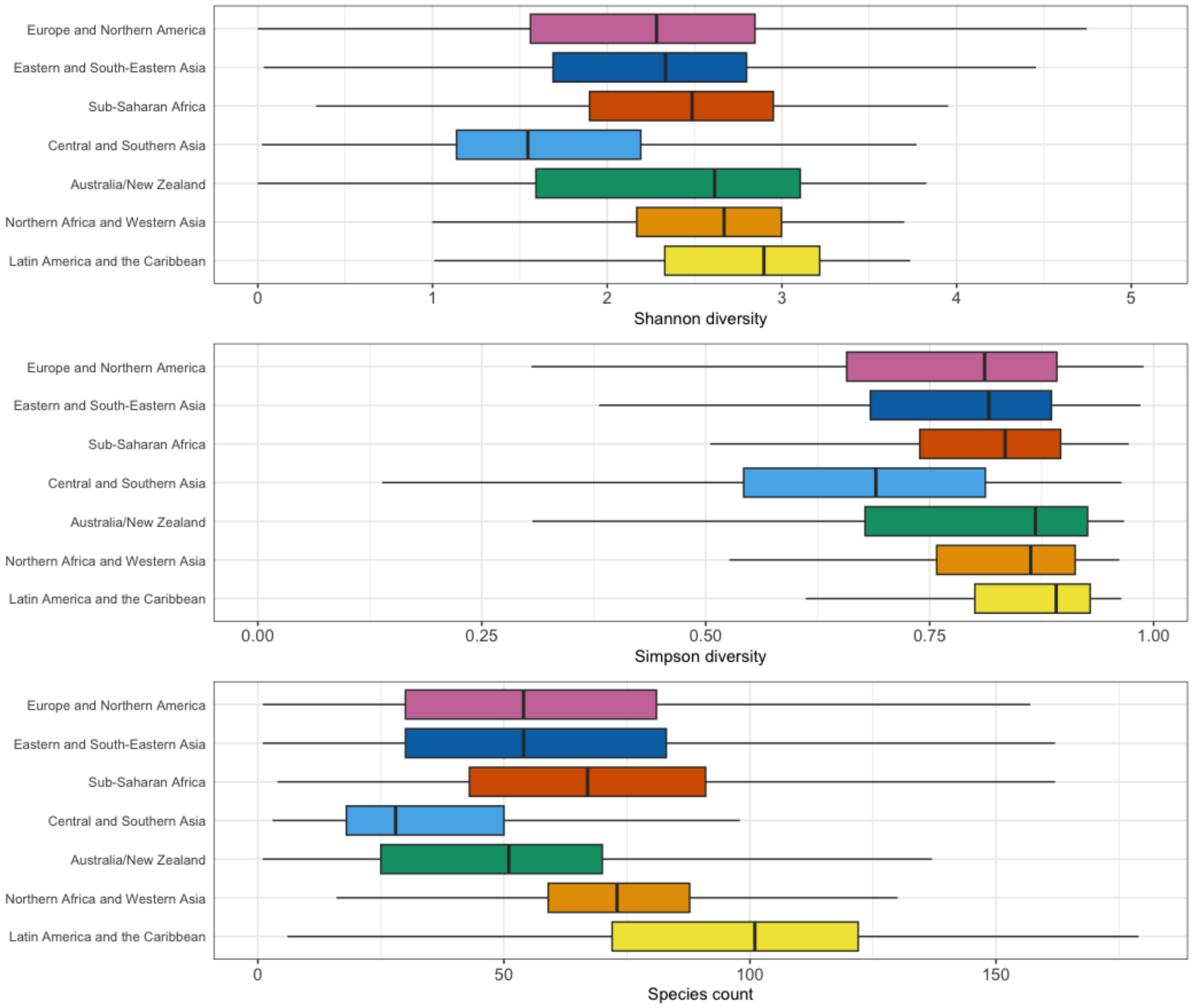


**Supplementary Figure 4. Diversity across world regions.** Similarly to **Figure 2C**, each panel illustrates the alpha diversity found in samples from each region, using different indices. The x-axis indicates the score, and the y-axis lists each region. The bold line within each box indicates the median, and the left and right box boundaries represent the first and third quartile respectively. The whiskers extending from each box indicates 1.5 times the interquartile range of samples from that region. The top panel uses the Shannon Index to measure alpha diversity in each sample; the middle panel uses Simpson's Diversity Index, and the bottom panel uses the "species count" as calculated by the vegan R package.

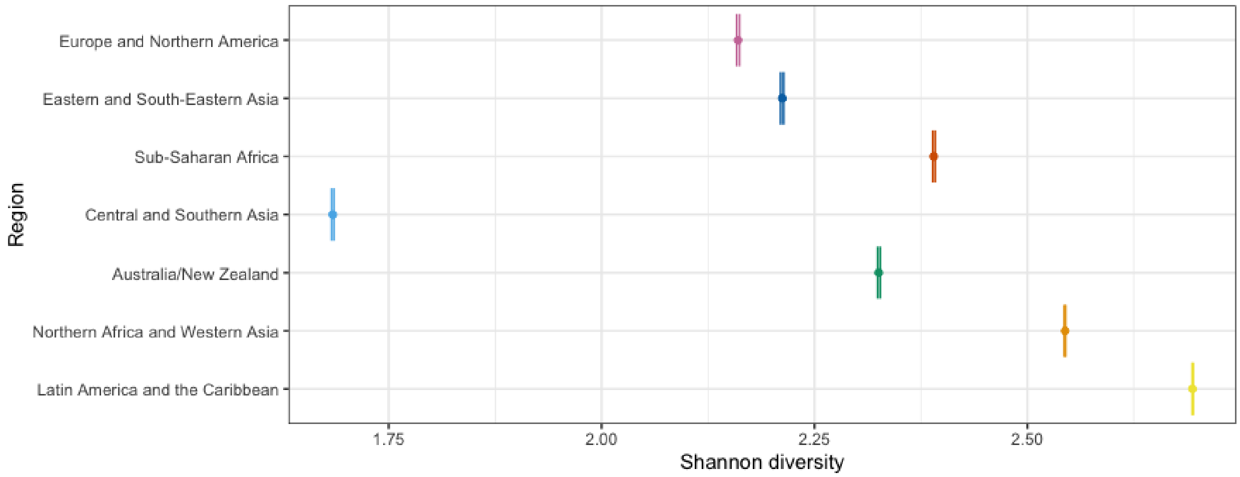


**Supplementary Figure 5. Rarefaction diversity estimates.** The x-axis indicates Shannon diversity, and the y-axis lists all regions evaluated in our rarefaction analysis. Each point indicates the mean alpha diversity calculated over 1000 iterations (see **Methods**). The error bars indicate the 95 percent confidence interval, calculated using the R "t.test" function.

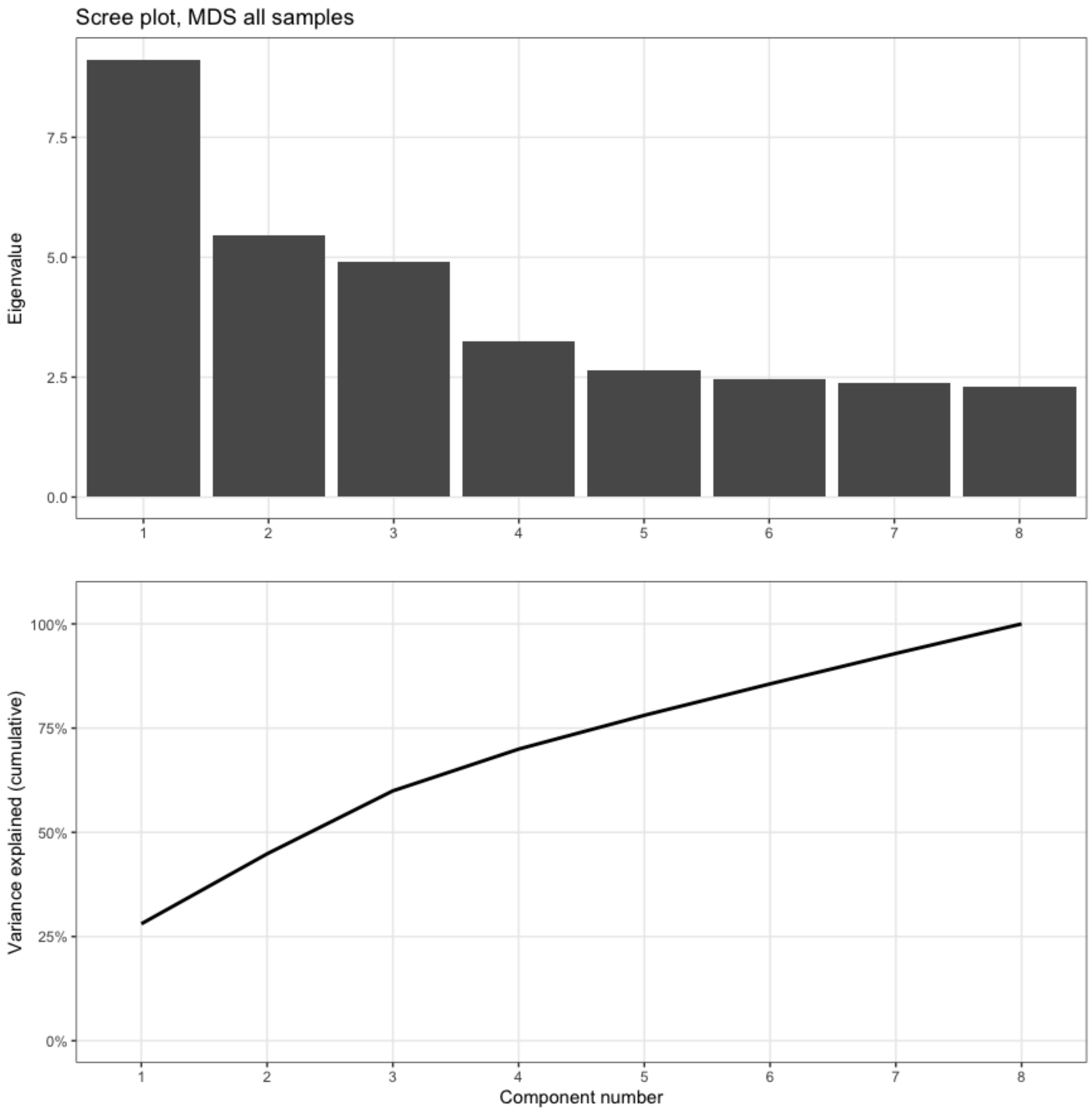


**Supplementary Figure 6. Scree plot for ordination analysis.** This plot illustrates the importance of all eight axes calculated during the ordination analysis illustrated in **Figure 2E–F**. The x-axis of both panels indicates the extracted components. In the top panel, the y-axis indicates the eigenvalue assigned to each component. These values are used to calculate the variance explained by each component individually; their cumulative total is illustrated in the bottom panel.

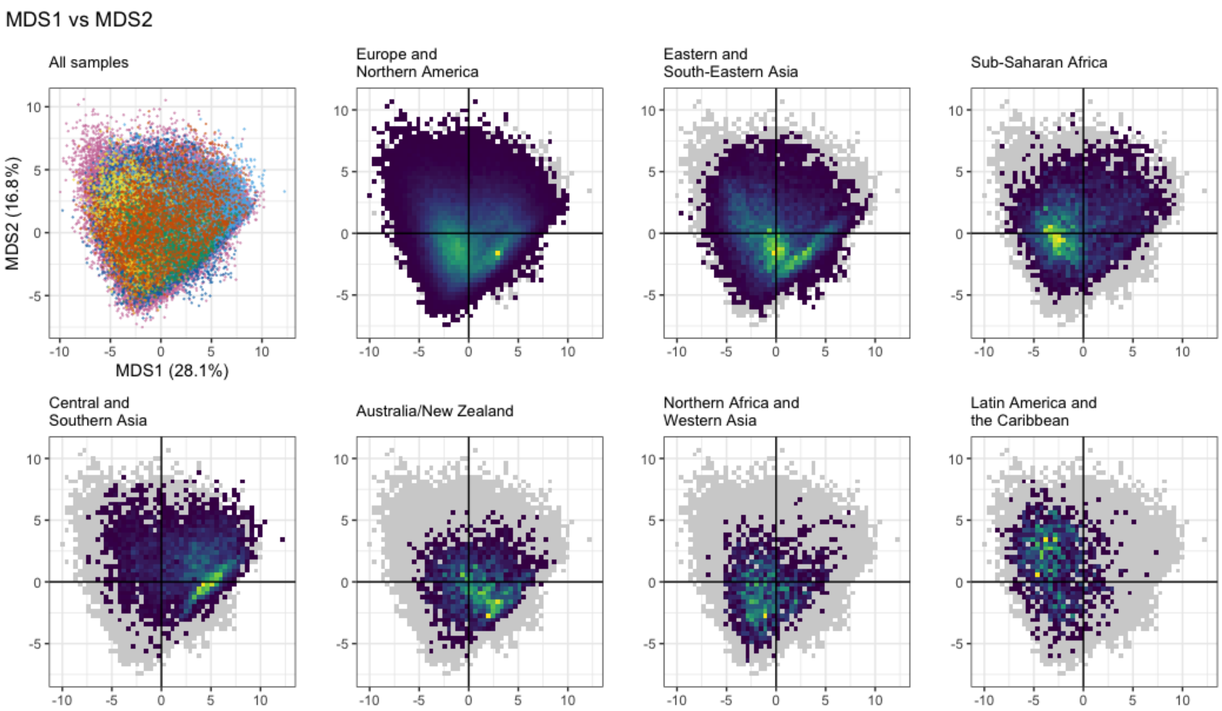


**Supplementary Figure 7. Alternate ordination plot.** The results of a principal coordinates analysis of samples from all regions. This figure is arranged in the same way as **Figure 2E**, but using a higher resolution (a 50-by-50 grid instead of 30-by-30). This figure plots patterns in the first and second MDS axes. The top-left plot is a scatter plot in which each point is a single sample; the color indicates the sample's region, using the scheme described in **Figure 2A**. The seven other plots use the same axes as the scatter plot, but each includes only samples from a single region. These plots use a heatmap design rather than a scatter plot to help evaluate areas with many overlapping points—yellow areas indicate portions of the space with a higher concentration of samples, and dark blue areas indicate portions in which few (but not zero) samples are found. The gray shadow added to each plot indicates the area occupied by all points from all regions.

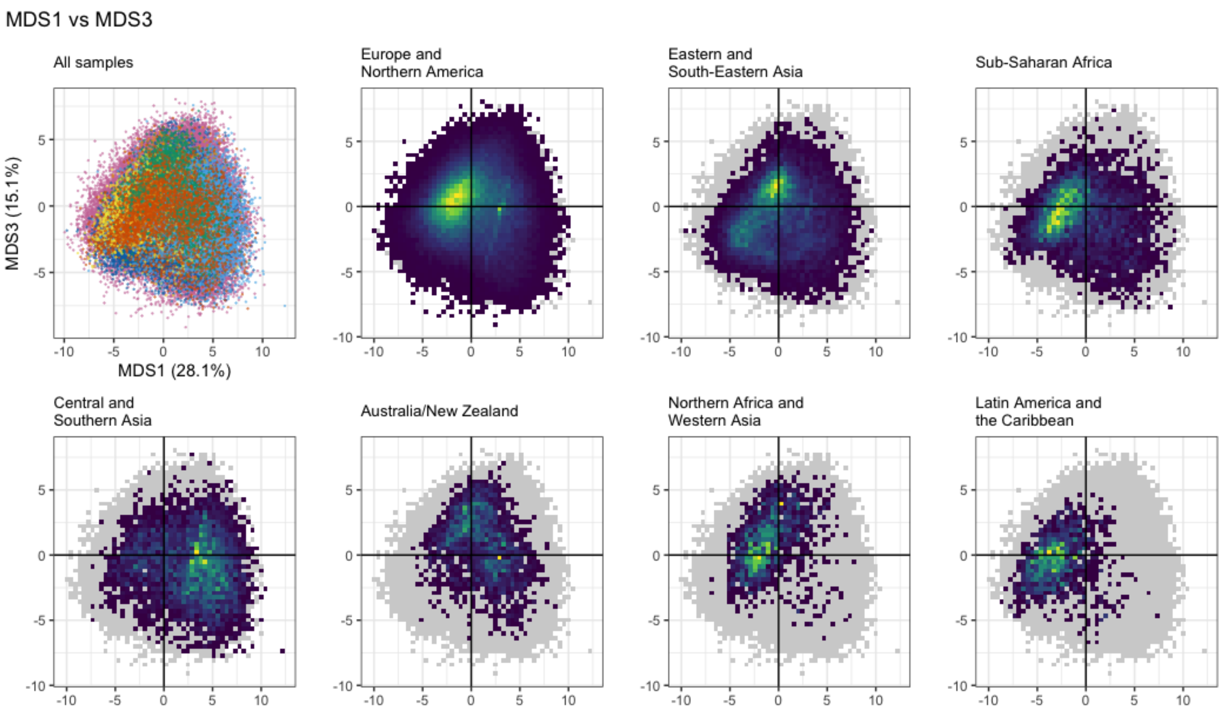


**Supplementary Figure 8. Alternate ordination plot.** Similar to Supplementary Figure 7, but showing axes MDS 1 vs MDS 3.

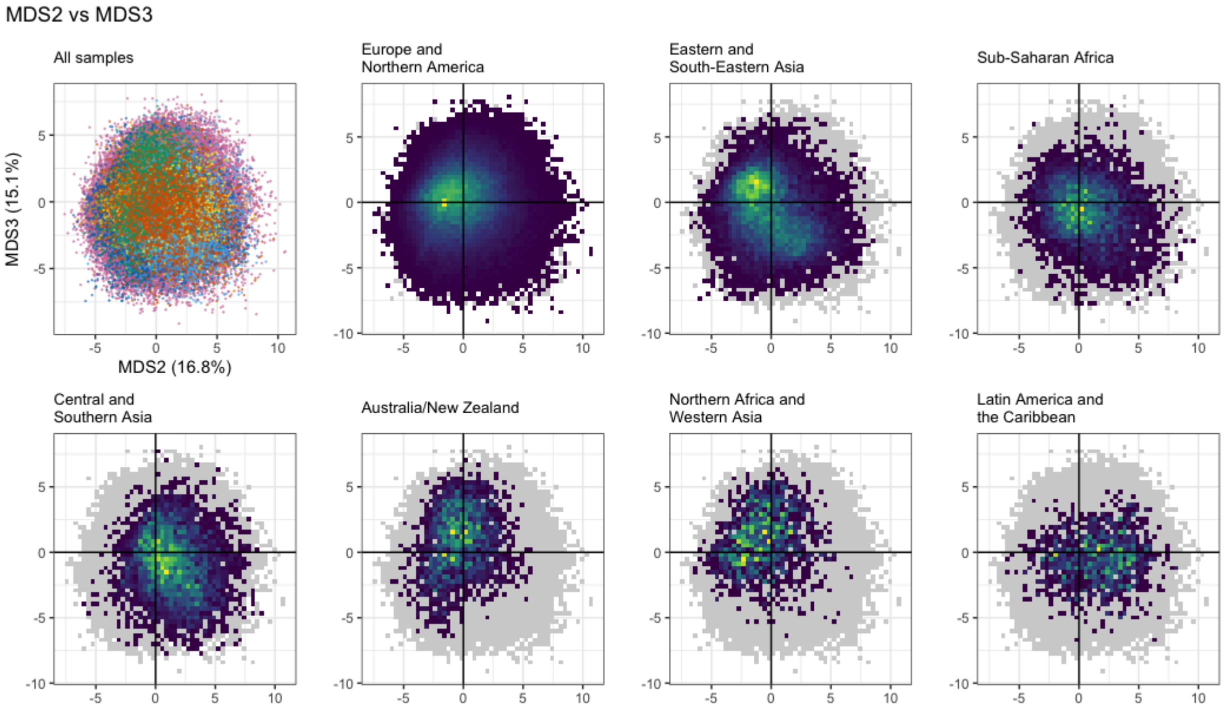


**Supplementary Figure 9. Alternate ordination plot.** Similar to Supplementary Figure 7, but showing axes MDS 2 vs MDS 3.

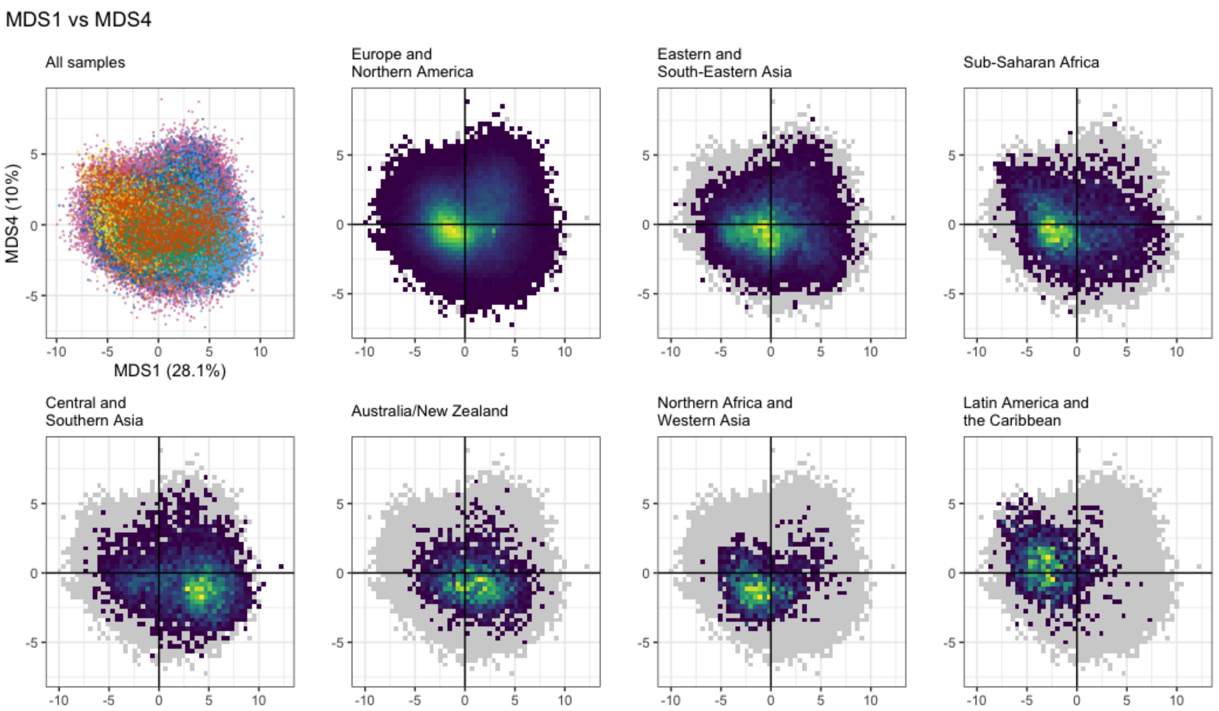


**Supplementary Figure 10. Alternate ordination plot.** Similar to Supplementary Figure 7, but showing axes MDS 1 vs MDS 4.

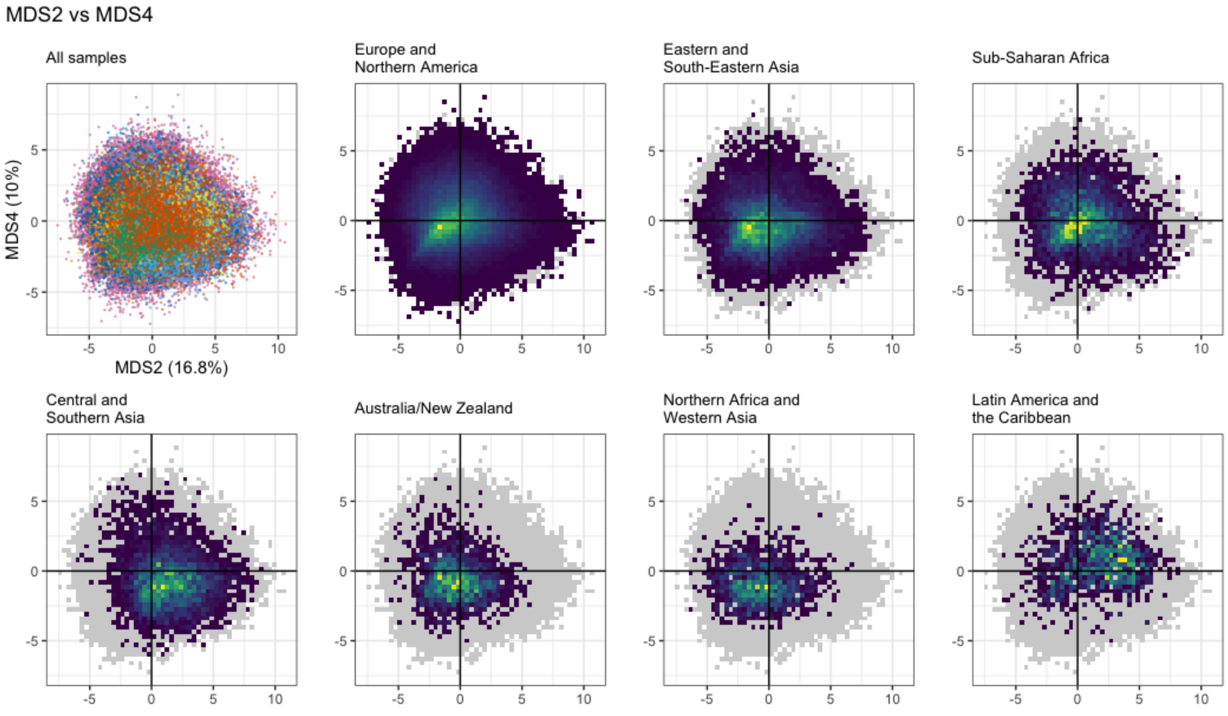


**Supplementary Figure 11. Alternate ordination plot.** Similar to Supplementary Figure 7, but showing axes MDS 2 vs MDS 4.

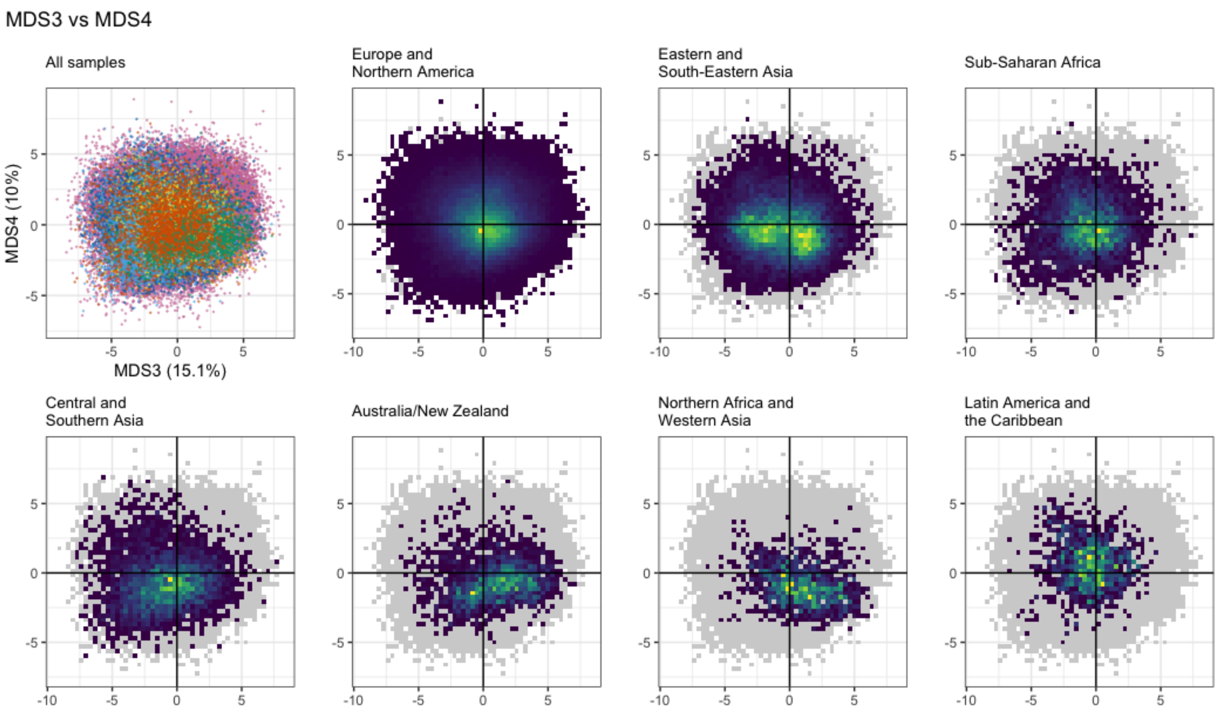


**Supplementary Figure 12. Alternate ordination plot.** Similar to Supplementary Figure 7, but showing axes MDS 3 vs MDS 4.

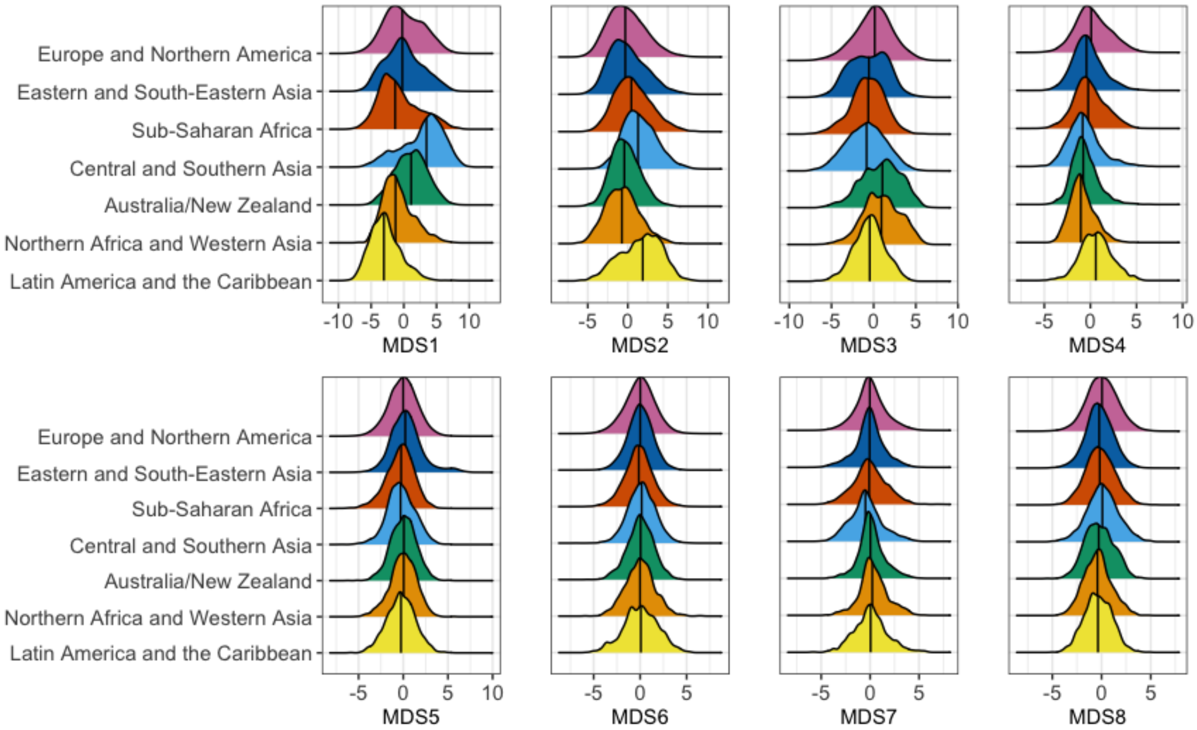


**Supplementary Figure 13. Ordination results.** A series of plots illustrating the results of a principal coordinates analysis of samples from all regions. This extends the data displayed in **Figure 2F** by showing factors MDS5 through MDS8. Each panel illustrates a single factor; the x-axis indicates the value of that factor, and the y-axis indicates the relative frequency of the value in the given region.

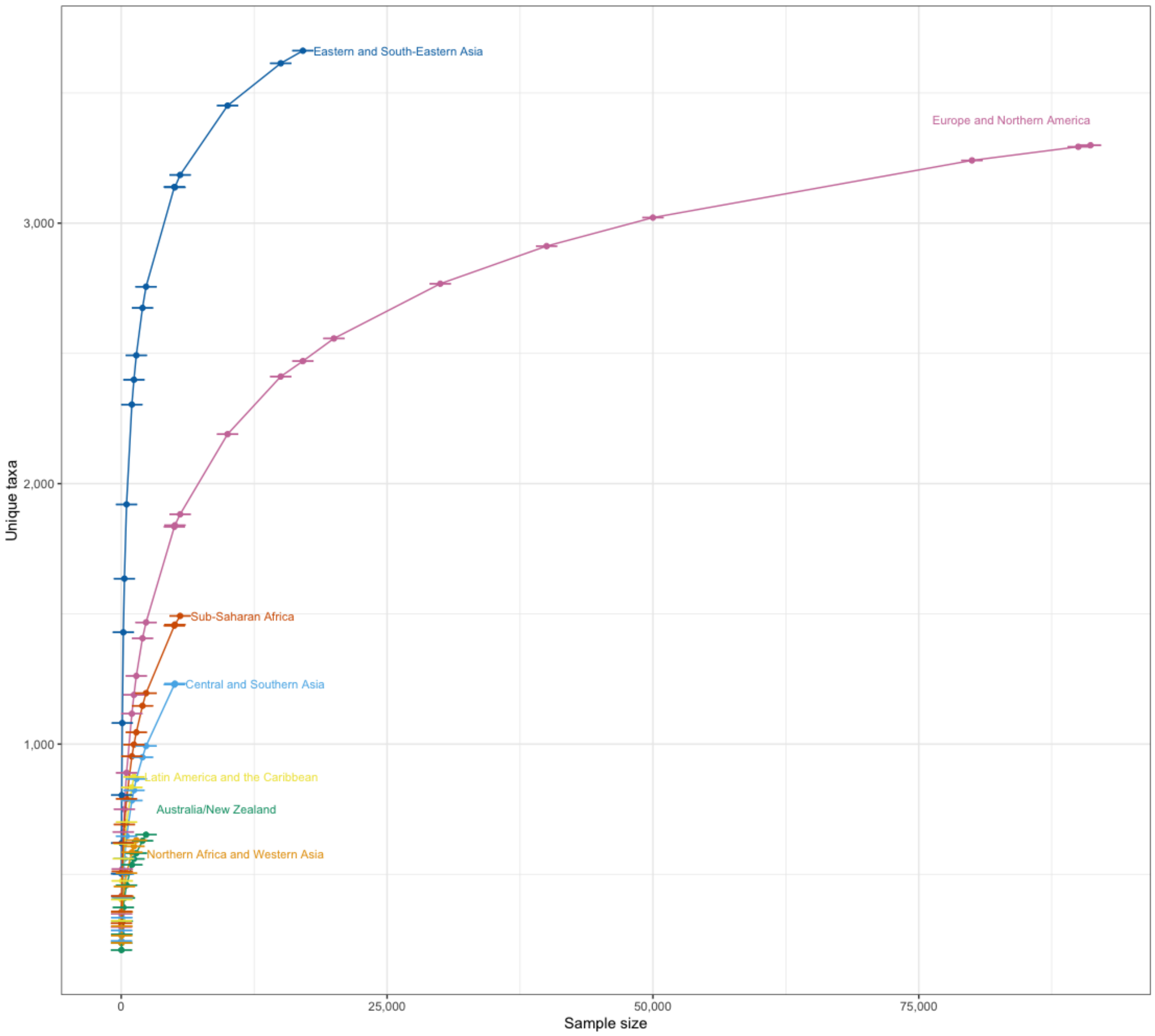


**Supplementary Figure 14: Region level discovery curve with 95% confidence intervals.** This plot reflects the same data as is presented in Figure 3A, with the addition of a 95% confidence interval of the unique taxa discovered at each sample size for each region. (Please note that the confidence intervals are illustrated vertically—the horizontal lines indicate the upper and lower bounds of the range, but the ranges are small enough that they appear very close together.)

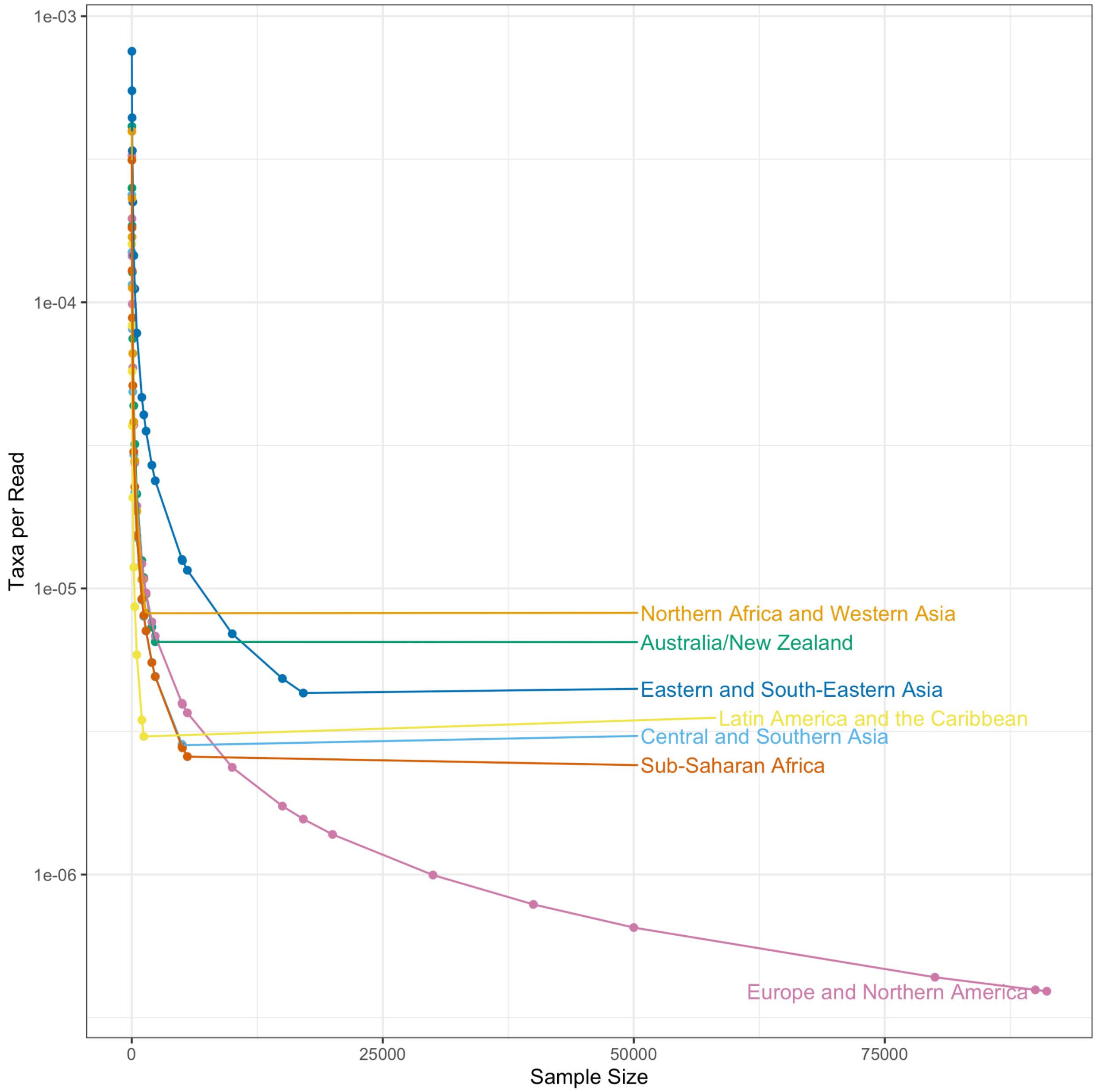


**Supplementary Figure 15: Region level discovery curve with number of samples on the x-axis and the number of taxa per read on the y-axis.** Similar to the inset in **Figure 3A**, we sampled increasing numbers of microbiomes from each world region. For each sample size (indicated on the x-axis), we calculated the number of unique taxa observed, and the number of reads present in each sample. We then calculated the mean Taxa per Read (indicated on the y-axis) by dividing the mean number of Unique Taxa by mean read count for each sample size.

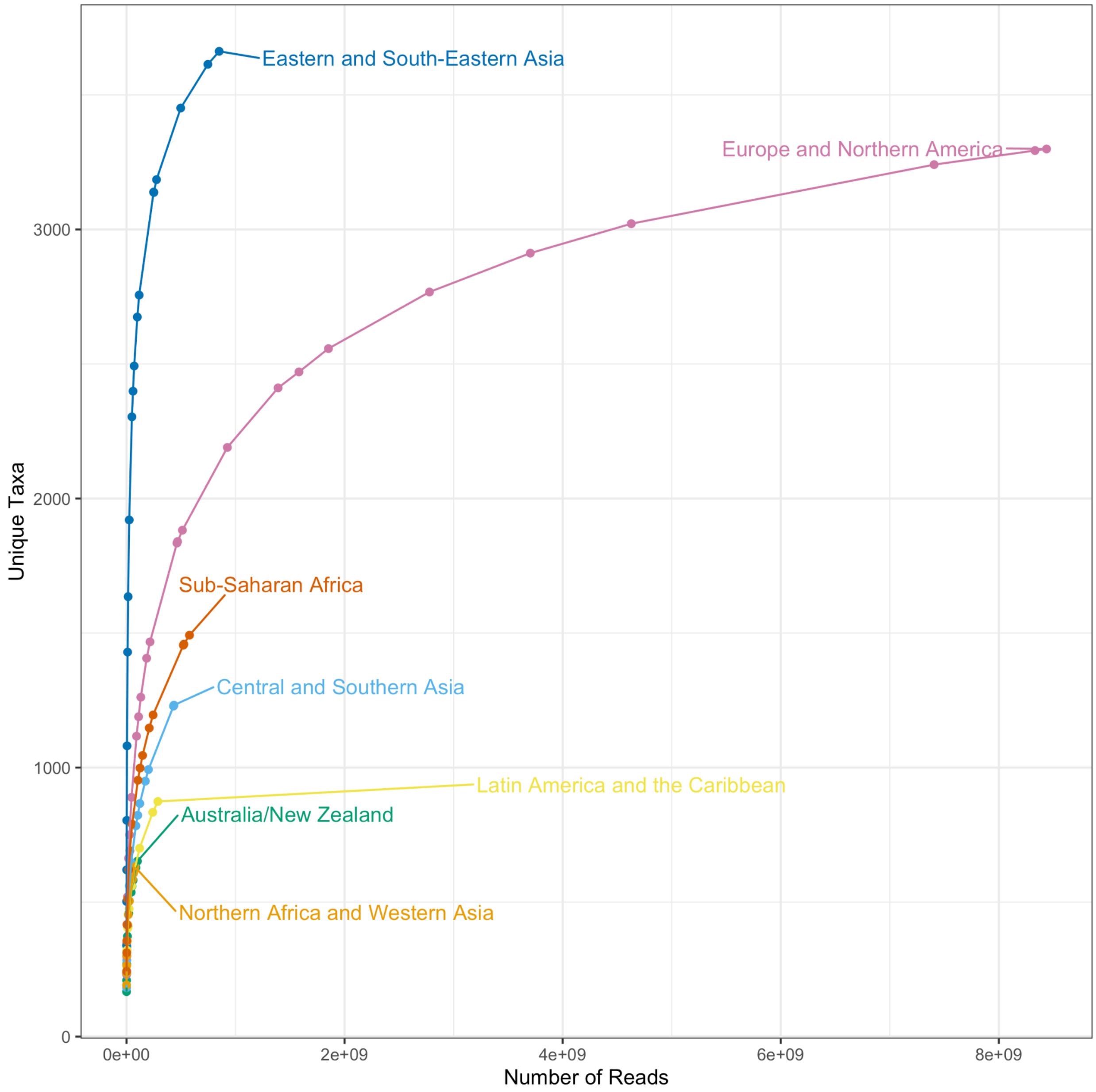


**Supplementary Figure 16: Region-level discovery curve plotting the number of taxa found as the number of reads increases.** Using the same sampling scheme as **Figure 3A**, we sampled increasing numbers of microbiomes from each geographic region. For each sample size, we calculated the mean number of reads in the selected samples (indicated on the x-axis), and the mean number of unique taxa observed in the selected samples (indicated on the y-axis).

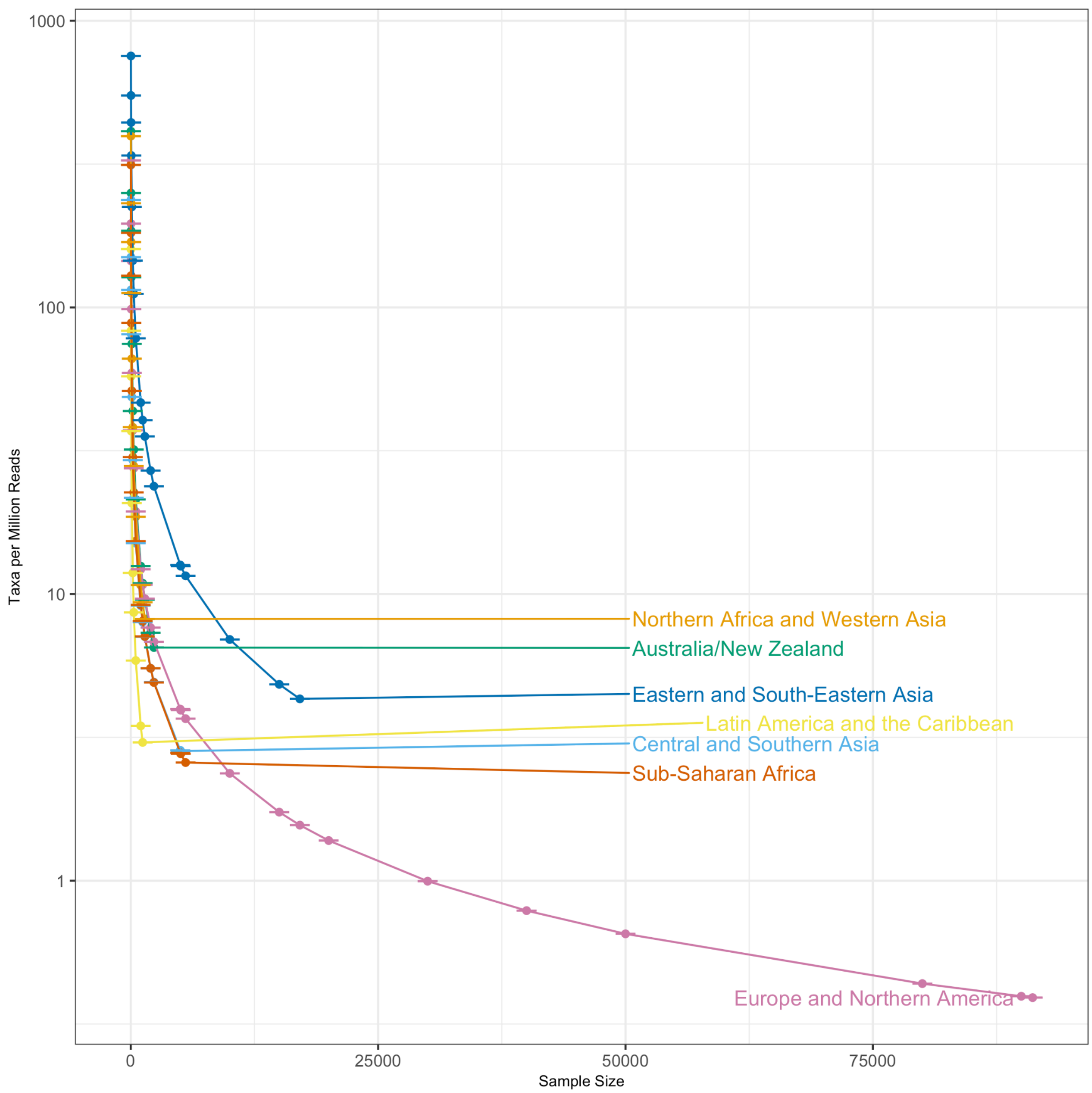


**Supplementary Figure 17: Region level discovery curve.** Similar to the inset in Figure 3A, with error bars showing the 95% confidence interval. The confidence interval was calculated by bootstrapping the mean Taxa per Million Reads 1000 times.

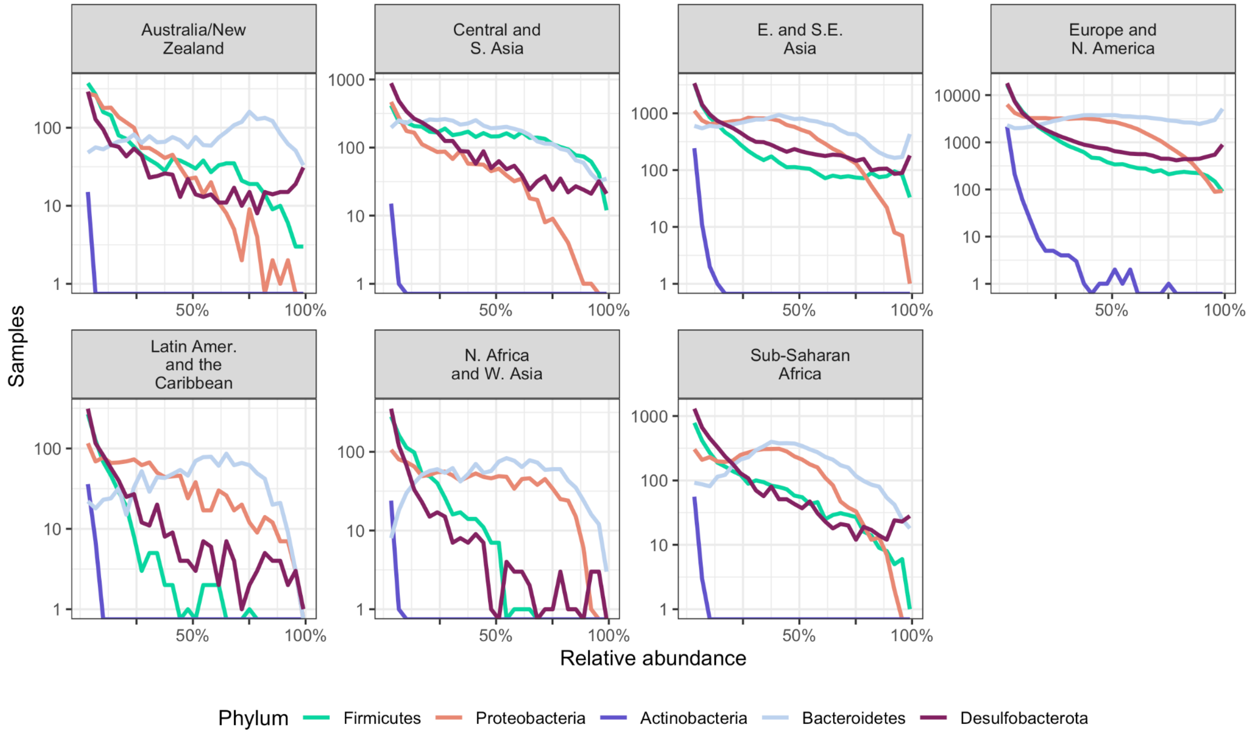


**Supplementary Figure 18: Histograms showing the relative abundance of different taxa across sample sizes, with individual y-axes for each region.** Similar to **Figure 3B**, these histograms illustrate the distribution of the relative abundance of the most prevalent phyla in the compendium. Each panel visualizes all samples from a single world region, with number of samples on the y-axis and relative abundance on the x-axis. The individual y-axes for each panel emphasize differences in sample size and microbiome composition between world regions.

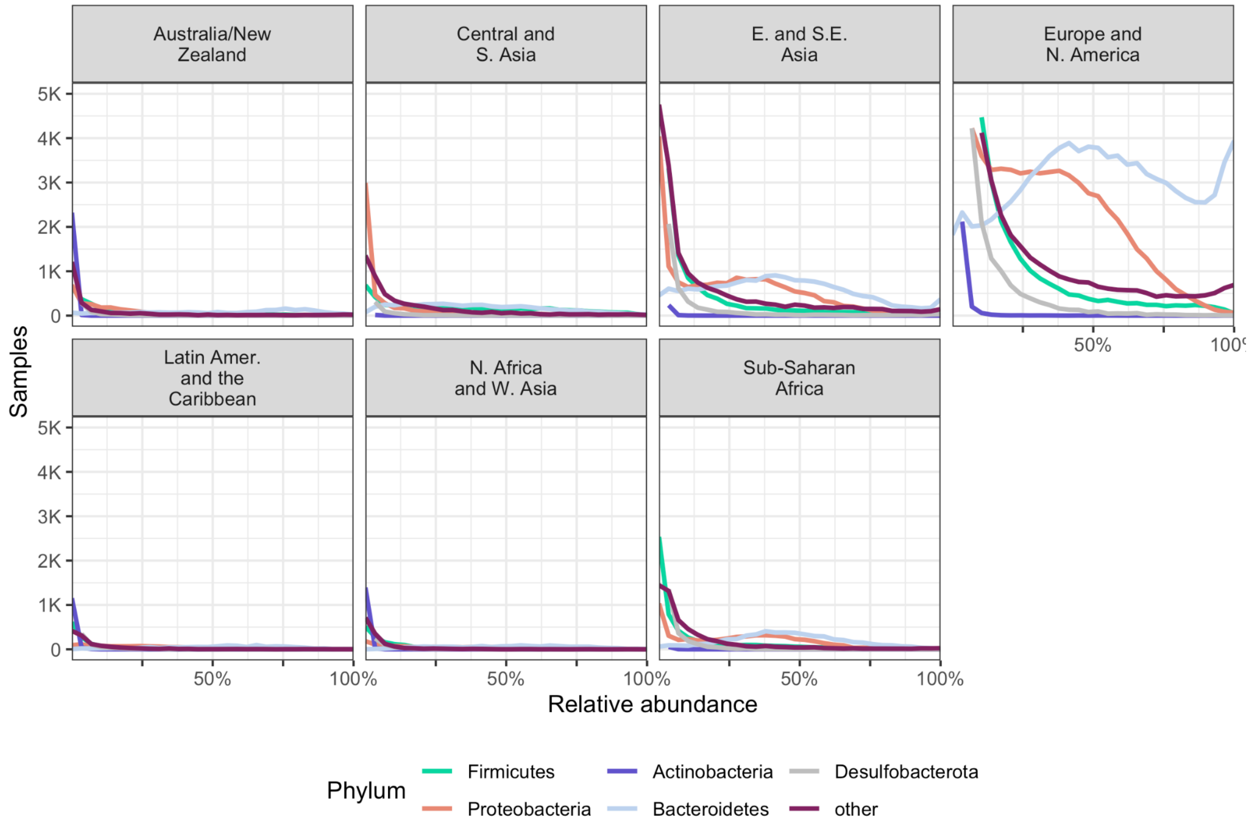


**Supplementary Figure 19: Histograms showing the relative abundance of different taxa across sample sizes, with a linear y-axis.** Similar to **Figure 3B**, these histograms illustrate the distribution of the relative abundance of the most prevalent phyla in the compendium. Each panel visualizes all samples from a single world region, with number of samples on the y-axis and relative abundance on the x-axis. The linear y-axis emphasizes the difference in sample sizes between world regions.

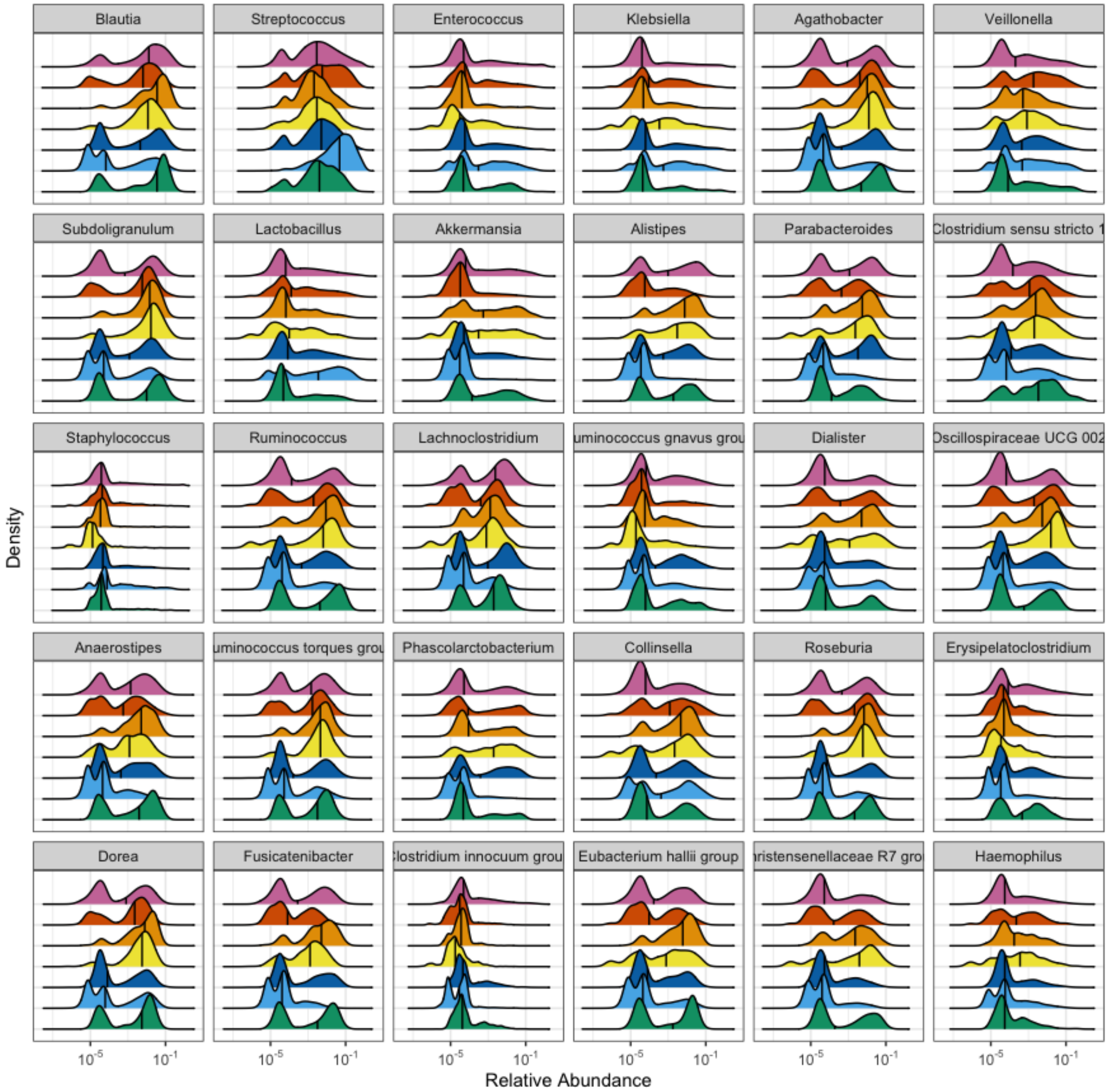


**Supplementary Figure 20: Relative abundance of differentially abundant taxa**. Similar to **Figure 4C**, each panel illustrates the relative abundance of a single taxon. Shown here are taxa 6-35 when ranked by average relative abundance. Each colored area in a panel indicates the distribution from a single world region. The x-axis indicates (log 10) relative abundance of the specified genus, and the y-axis indicates the relative frequency with which that abundance is observed in the specified region. Black vertical lines indicate the median.

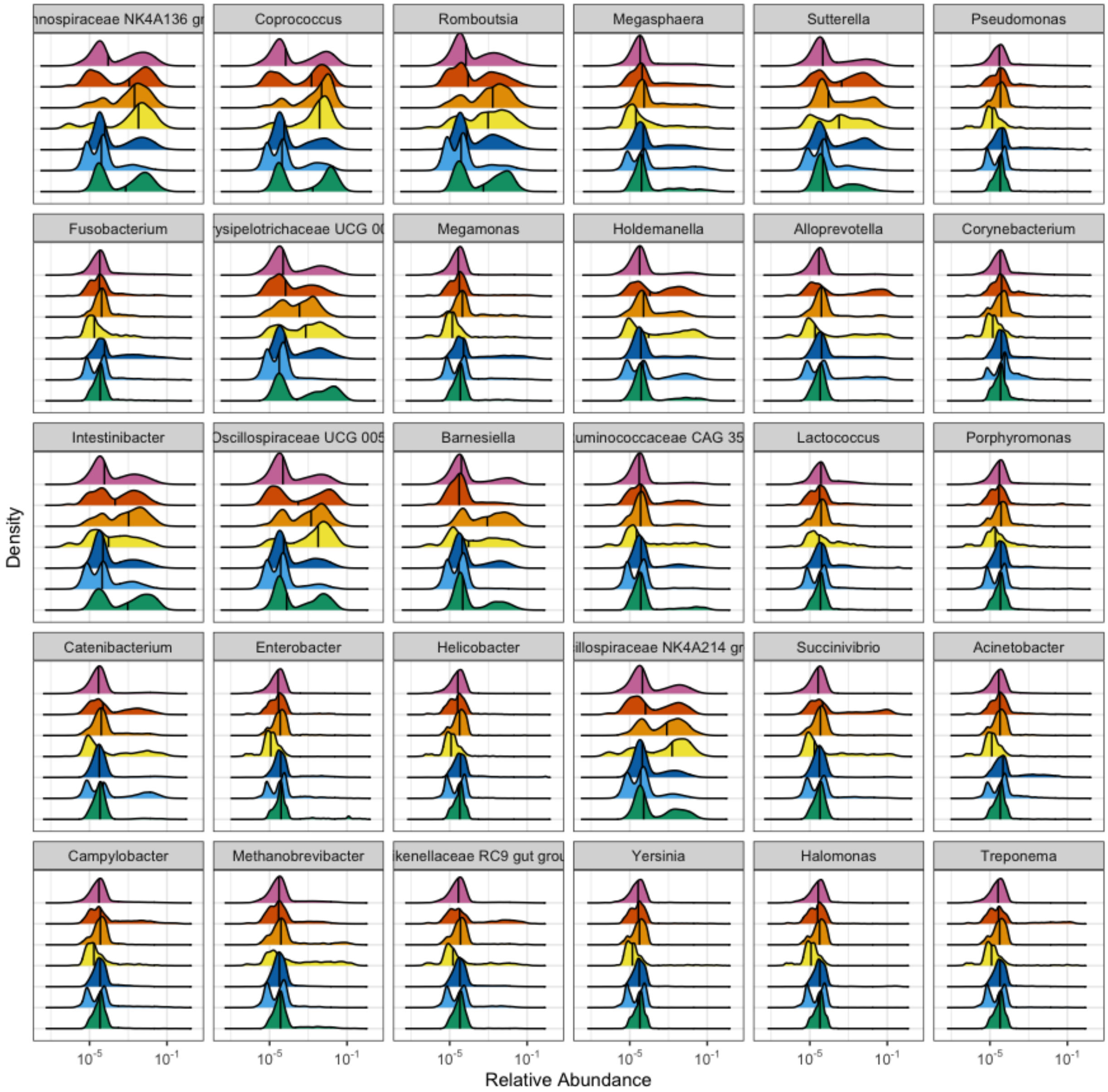


**Supplementary Figure 21: Relative abundance of differentially abundant taxa**. Similar to **Figure 4C**, each panel illustrates the relative abundance of a single taxon. Shown here are taxa 36-65 when ranked by average relative abundance. Each colored area in a panel indicates the distribution from a single world region. The x-axis indicates (log 10) relative abundance of the specified genus, and the y-axis indicates the relative frequency with which that abundance is observed in the specified region. Black vertical lines indicate the median.

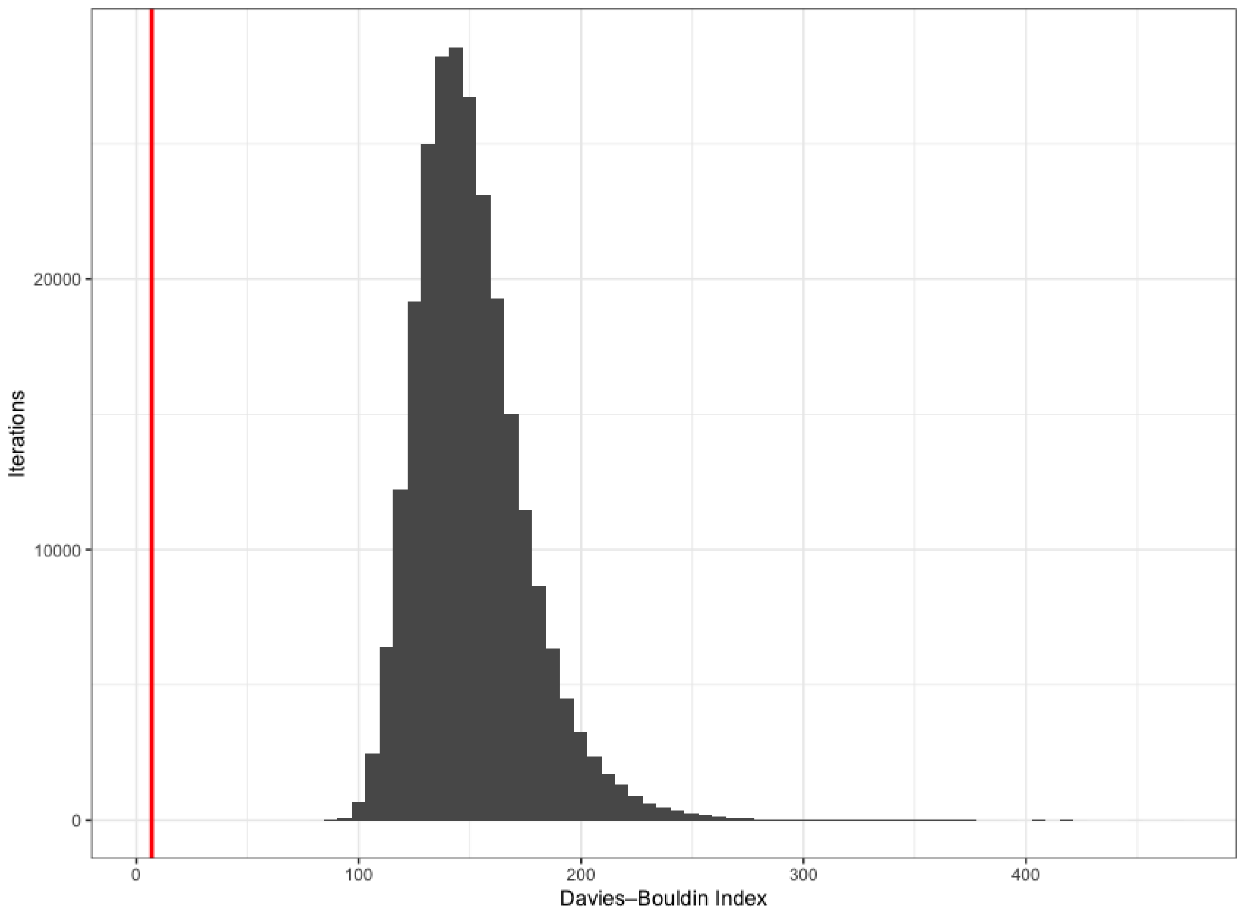


**Supplementary Figure 22. Cluster strength analysis.** A histogram illustrating the results of a bootstrap analysis evaluating the strength of the regional clusters formed in the ordination space. Each iteration (total 250,000) shuffled the region labels attached to all samples and generated a single score (Davies–Bouldin Index) for the iteration. The x-axis indicates the score, and the y-axis indicates how many iterations had a score in that bin. The red line indicates the score (6.93) for clusters defined by the real regions.
